# ModuloStat: An Internet of Things’ Path to Continuous Cultures in Mini-Bioreactors

**DOI:** 10.1101/2025.08.08.669317

**Authors:** Cyprien Guérin, Etienne Dervyn, Joséphine Véron, Ira Tanneur, Sandra Dérozier, Magali Calabre, Elena Bidnenko, Matthieu Jules, Pierre Nicolas

## Abstract

In continuous culture, a population of microorganisms is propagated in a stable environment over many generations. This is particularly relevant for experimental evolution and metabolic studies. However, continuous culture protocols are difficult to implement, so they are not commonly used in microbiology laboratories. Here, we present the ModuloStat, a modular, open-source framework that facilitates continuous culture in mini-bioreactors. The ModuloStat system is grounded on digital fabrication tools easily accessible in FabLabs and programmable electronics. Maintaining a culture is divided into tasks assigned to dedicated printed circuit boards with a microcontroller connected to a Wi-Fi network. According to Internet of Things principles, each board operates a set of sensors and actuators autonomously and can receive and send information. The boards are stacked to implement complex behaviors and can be modified to accommodate new features. A thermoregulated box holds the components and can be placed on a laboratory bench or transported under a sterile hood for inoculation. Sterility is ensured by autoclaving, after assembly, all components that will come into contact with the culture medium. *In-situ* optical density monitoring combined with modularity and computer control enables many cultivation modes. Additionally, we present the construction of the *Bacillus subtilis* strain ZB designed for bioreactor culture that exhibits a zero-biofilm phenotype. To demonstrate the system’s versatility, we performed several experimental cultures with this model organism, including chemostat, turbidostat, medium swap, and a cascade of bioreactors.

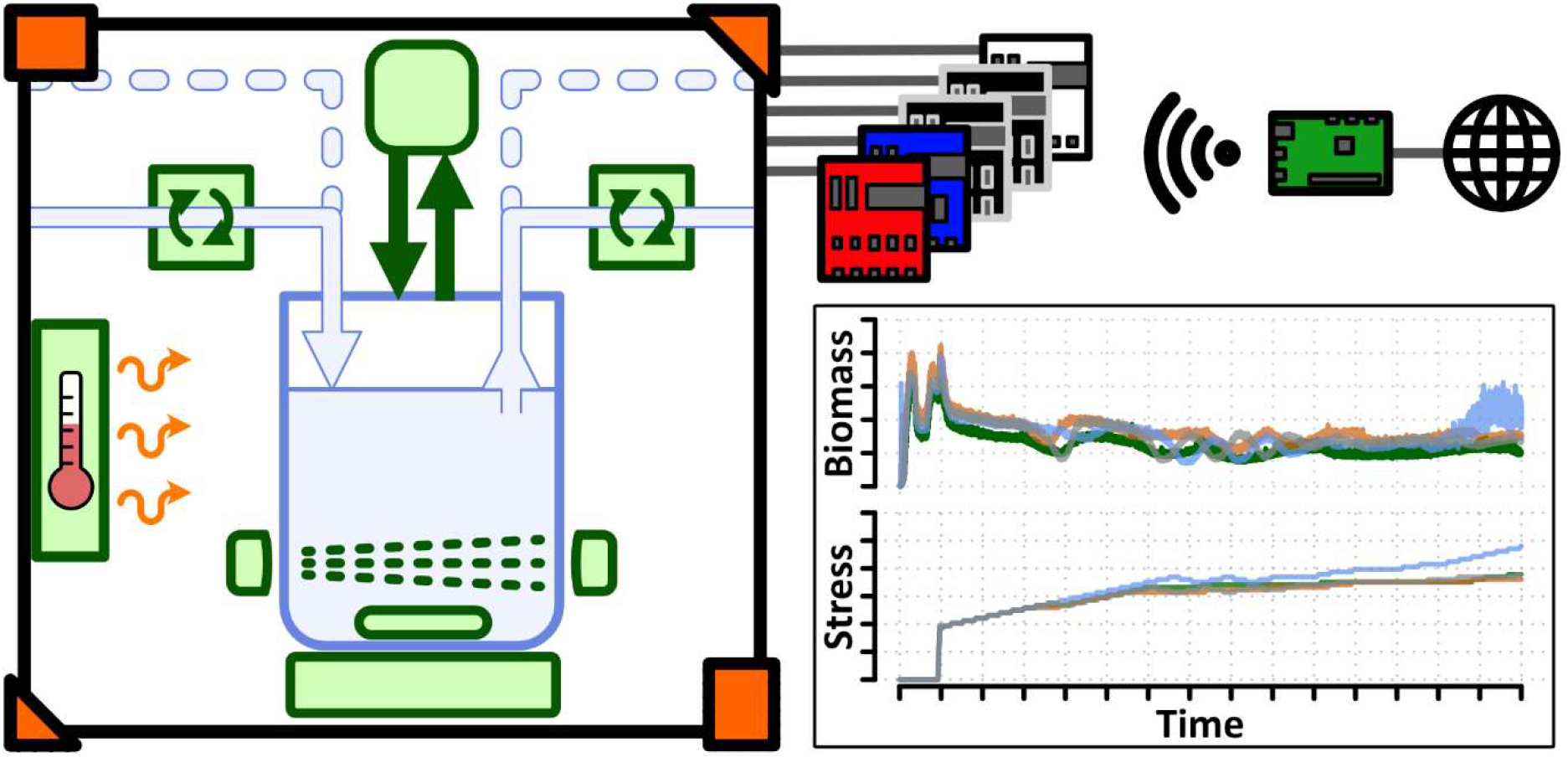

## INTRODUCTION

Within a single day, a liquid culture consisting of a rich medium inoculated with a few cells of a fast-growing bacterium passes through a succession of growth phases, each characterized by the exploitation of different resources. In this “batch” setup, saturation is reached at ≈10^9^ bacteria per mL after less than ≈40 generations for one litter of culture; doubling or halving the volume adds or removes only a single generation. A specialized apparatus is required to achieve continuous growth over a large number of generations in the laboratory. The simplest mode of continuous culture is the chemostat, which maintains a constant-volume culture fed by a continuous inflow of fresh medium, with volume controlled by an overflow (^1,2^). Mimicking a simple lake, the chemostat allows stable physiological conditions and provides the ideal environment to study metabolism and competition in a steady state (^3^). The chemostat also stands as a paradigm for more complex natural or artificial open systems such as some large bioreactors used in wastewater treatment or industrial fermentation and bioproduction processes (^4,5^). In addition, continuous cultures are of foremost interest for adaptive laboratory evolution (^6–8^). For all these reasons, continuous cultures have been the focus of the development of sophisticated mathematical models at the interface between physiology, ecology and evolution (^9,10^). Nevertheless, despite their attractiveness and early introduction, most microbiology laboratories do not routinely use these systems. In fact, the chemostat and its more complex derivatives such as turbidostat (^11^), gradostat (^12^), or morbidostat (^8^), remain difficult to implement.

Another significant challenge to the practical application of continuous culture is that microbes can swiftly circumvent dilution by forming biofilms or aggregates. This limits the prolonged use of systems lacking specific mechanisms to prevent the proliferation of dilution-resistant variants (^13^). One solution is to stop the culture after one day and restart it from a sample (^14,15^). Other options consist of manually (^16^) or semi-automatically (^17^) replacing bioreactor vessels during the culture. A more complex yet more satisfying approach is to automate the periodic transfer of the entire culture between two vessels, as well as the intercalated cleaning of the empty vessel (^18^).

The modern era of continuous culture systems began with the manual assembly of commercially available materials in order to perform adaptive laboratory evolution (^8,19^). Shortly after, more do-it-yourself bioreactor implementations for physiological, evolutionary, and synthetic biological approaches emerged (^17,20–23^), including reimplementations of the popular morbidostat design (^15,24^). These implementations leveraged digital fabrication and programmable microcontrollers or single-board microcomputers. The proposed systems cover a broad range of bioreactor volumes (from 12 to 800 mL) and incorporate various sensors and actuators to control growth (see Table 1 for a summary). These systems were developed with technical choices that optimize specific use cases. For example, air-pushed overflow minimizes the number of pumps, enabling compact, parallel mono-bioreactor systems (^8,20^), while highly integrated printed circuit boards (PCBs) streamline production and assembly (^17,24^). However, these central technical choices inherently limit the versatility of the proposed systems. Adapting them to other experimental designs may require significant modifications or nearly complete refactoring. Nevertheless, the authors of these recent systems emphasize sharing their designs, from parts assembly to dedicated control software, including electronic circuits and 3D models to print. Different levels of sharing are distinguished in Table 1.

**Table 1:**
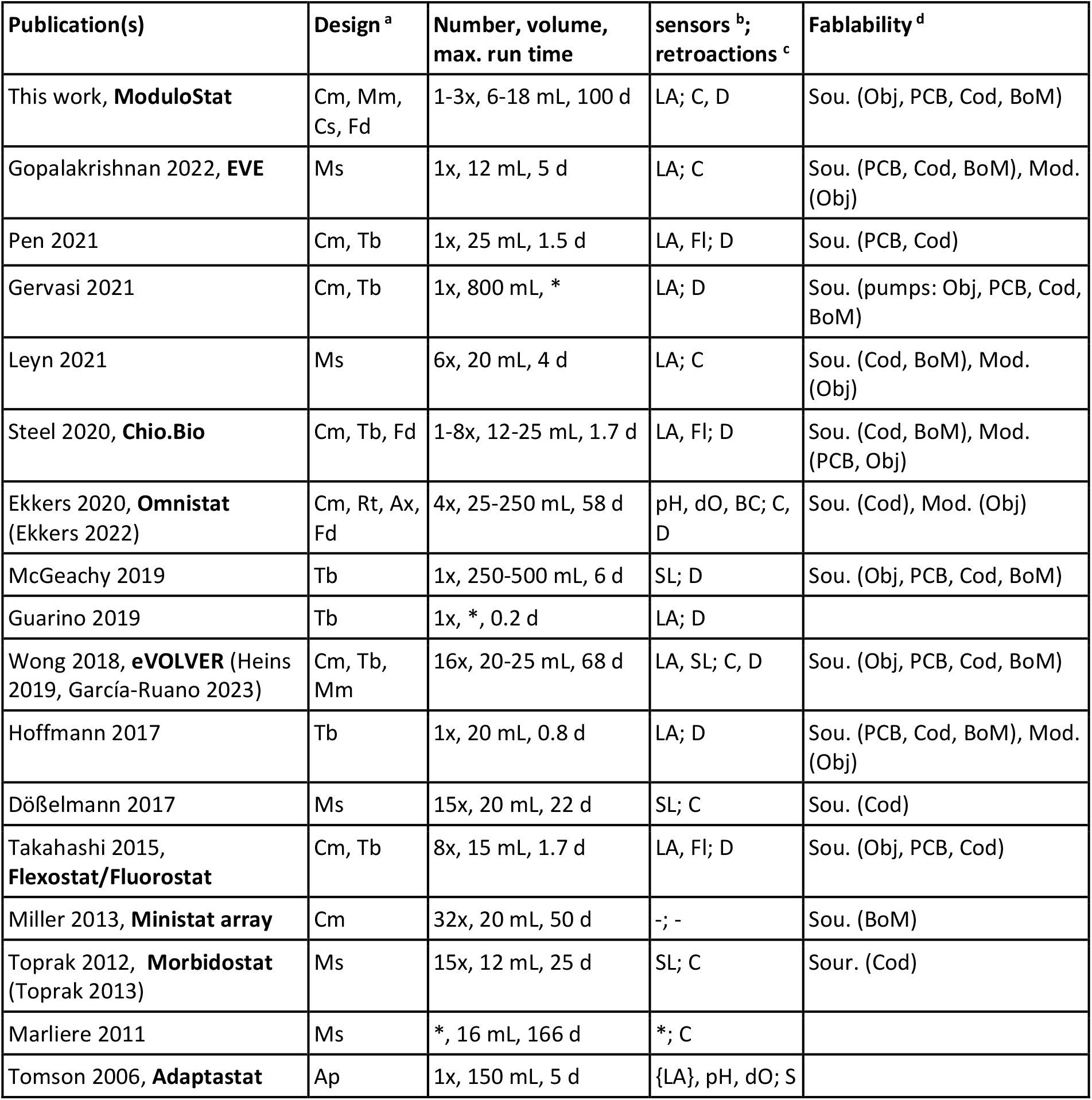
Cultures characteristics of already published continuous culture systems. ^a^ Cm (chemostat), Tb (turbidostat), Mm (medium mix), Ms (medium swap), Cs (cascade), Rt (retentostat), Ax (auxostat), Ap (adaptastat), Fd (flexible design, other possible applications mentioned in paper but not presented). ^b^ - (none), LA (light absorbance), SL (scattered light), Fl (fluorescence) pH (potential of hydrogen), dO (dissolved oxygen), BC (biomass capacitance), not *in-situ* measures between brackets. ^c^ - (none), C (chemical composition by medium mix or swap, or chemical injection), D (dilution rate), S (stirring speed). ^d^ Sou. (sources available allowing easy modifications) or Mod. (models or technical drawing available allowing reproduction). Listed for: Obj (objects), PCB (electronic boards), Cod (codes), BoM (bill of material), AsI (assembly instructions). * Information not available in publication, supplementary information or associated online data.

These developments in bioreactor systems for continuous culture are part of broader technological and societal trends. Easy access to digital fabrication, facilitated by desktop 3D printers and tools available in nearby FabLabs, promotes the development and spread of well-documented projects that can be reproduced virtually anywhere around the globe (^25^). It is hoped that this will give citizens, including researchers, increased control over the factors that influence their daily environment, a concept known as empowerment (^26^). Programmable microcontrollers, such as Arduino (https://www.arduino.cc/) and ESP32 (https://www.espressif.com/en/products/socs/esp32); single-board computers, such as Raspberry Pi (https://www.raspberrypi.com/); and accessible prepackaged electronics, such as Adafruit products (https://www.adafruit.com/), have played a major role in facilitating interactions between computers and the real world. They enable the development of devices equipped with sensors and processing capabilities that can connect to and exchange data with other devices and systems via communication networks. This leads to the rapid growth of an extension of the Internet known as the Internet of Things (IoT); an evolution that is accompanied by the standardization of computer network communications for data gathering and transfer. Examples include domestic applications in “smart” homes, but they are also starting to penetrate laboratory equipment (^27^).

In this work, we introduce the ModuloStat, an open-source framework designed to facilitate the modular fabrication, assembly, and automated control of continuous culture in mini-bioreactors based on digital fabrication and IoT. First, we present and discuss the specifications and design choices of this framework. Next, we present and experimentally validate its central components, including pumps, optical density (OD) sensors, and thermal regulation. Concurrently, we describe the development of a new strain of the important model bacterium *Bacillus subtilis* that is genetically engineered to produce a zero-biofilm phenotype; this strain enables prolonged continuous culture in mini-bioreactors. Lastly, we demonstrate the application of the framework by implementing mini-bioreactor systems with various topologies, including simple chemostats, an OD-driven medium-mixing protocol for adaptive laboratory evolution to increasing ethanol stress, and continuous culture of a bacteriophage in a cascade of bioreactors.

## RESULTS AND DISCUSSION

### System specifications

We designed our system around typical culture volumes of approximately 7 mL, which is slightly lower than that of all previously described continuous culture systems to which we compare (Table 1). In a continuous culture, a steady state is reached when the growth rate, which is proportional to the inverse of generation time, matches the dilution rate (Supplementary Text). To set a chosen dilution rate, the flow of medium in the bioreactor is proportional to its volume, with several important implications in terms of practical use: medium consumption and waste production increase proportionally with the volume and that the time needed to collect a biological sample of a given size from the outflow decreases as the inverse of the volume. Small volumes achieve an interesting trade-off, particularly when the focus is on long-term evolution (^19,20^). With a 7 mL volume, a 10 L bottle of fresh medium can sustain ≈2,061 generations of a culture in steady state (Supplementary Text). This corresponds to approximately 43 days of experimental runtime if the generation time is 30 minutes, with an outflow that permits collecting a 1 mL sample in 6.25 minutes. These non-invasive samples are useful for creating a frozen record of the evolutionary trajectory, from which populations can be sequenced and single clones can be isolated.

Sterility is a key factor in long-term experiments. In our system, the passive component surrounding the fluidic circuit consists of an assembly of glassware, silicone tubing, and air filters which can be assembled, connected to a bottle of fresh medium, autoclaved, and installed in the system later, similar to the method described by Gopalakrishnan (^24^). After sterilization, dedicated sections of flexible tubing are fitted inside peristaltic pump heads without opening the circuit, thus avoiding any compromise to sterility (Figure 1). This eliminates the need for chemical sterilization of the connectors or larger portions of the circuit, as is required in some other systems (^17,23^; Table S1A). Our pumps are driven by low-voltage stepper motors that rotate in a series of small angular steps in response to user instructions, allowing for precise flow control (^27^). In contrast, other types of motors may not allow for direct command of the number of rotations (^24^).

**Figure 1:**
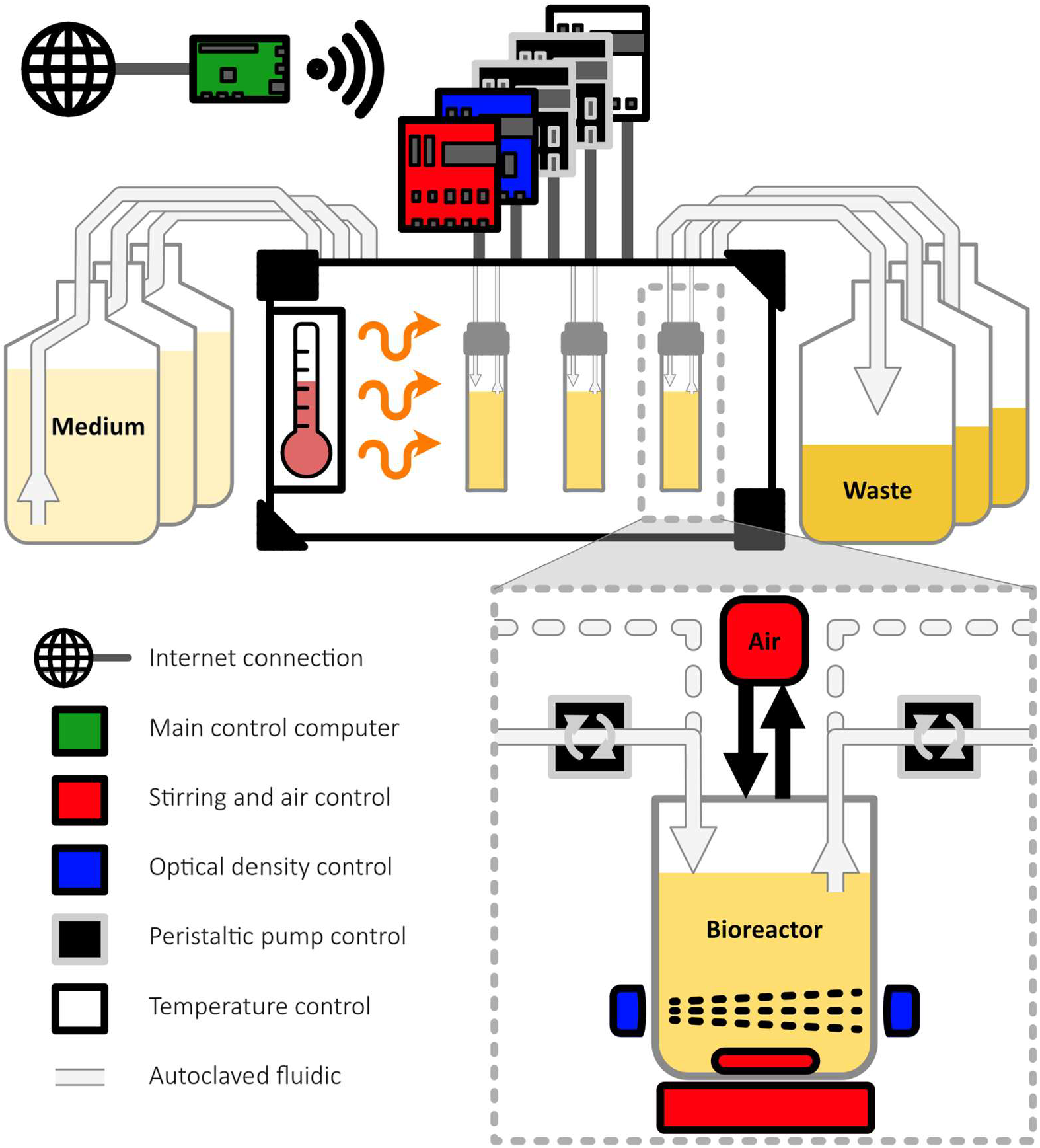
Schematic representation of the ModuloStat system and control. The main mini-computer (green) used for managing experiments is connected to the internet, allowing remote control. The boards (IoT devices) with different recipes (in red, blue, black, and white) communicate with the mini-computer via Wi-Fi and are connected to the actuators and sensors in the box, represented with the same color as the corresponding board. The autoclavable fluidic system (light gray) consists of mini bioreactors connected to feeding and waste bottles. The zoom on a mini-bioreactor shows optional additional fluidic inputs and outputs for more complex experimental designs.

Generation time heavily depends on temperature, and 37°C is a typical growth temperature for many fast-growing microorganisms. The easiest option for setting the temperature surrounding a continuous culture system apparatus in the laboratory is a heated room (^8^). However, a shared room is typically susceptible to temperature oscillation during working hours. High ambient temperature also imposes harsh conditions on the entire system, including the electronics and motors, as well as on human intervention. Alternatively, as presented in Table S1B autonomous temperature control allows precise parameters to be set for each experiment and can be achieved using a heated aluminum casing (^17^), a thermostated water bath (^28^), or by heating the air inside a small chamber (^15^). The ModuloStat framework uses the latter solution because it leaves the view to the culture, it raises the temperature of the medium in the tube on its way to the bioreactor, and it maintains the culture’s temperature if it has to be transferred between two bioreactors. However, it is less responsive if temperature variation is required during the experiment and it imposes the same temperature on all bioreactors in the same chamber. The chamber was designed under the form of a transportable box that can be moved under a sterile hood for culture inoculation (Figure 1). It can also be placed in a controlled environment for anaerobic conditions or for the use of dangerous substances. The heat-insulated box also limits light pollution, preventing interference with light sensors and physiological responses elicited by light. A transparent window with a lid allows for direct viewing of the culture when needed.

Light absorbance is the logarithm of the ratio of the power incident on a sample to the power transmitted through it. After division by optical path length, the result is the OD, which is commonly used in microbiology laboratories as a proxy to monitor changes in biomass concentration. *In-situ* measurement of light absorbance can be performed when bioreactors are small in volume by placing a light emission source and a receptor on the opposite sides of the bioreactor. This principle has been implemented in most of the systems proposed for continuous culture, although some systems instead use the scattered light fraction (see Table 1 and Table S1B). In these systems, the receptor is placed aside from the light emission beam, which may be advantageous when the amount of transmitted light is too low. Although 600 nm is a standard wavelength used in the laboratory, as about half of the systems already proposed we choose to measure OD in the near-infrared (i.e. above 700 nm). Then, OD measurements benefit from the virtual absence of indoor ambient infrared light, and the visible spectrum can be used for visual inspection of cultures and the surrounding system. Agitation is needed in a continuous system in order to homogenize the content of the bioreactor and it also speeds up gas-exchange at the air-liquid interface. Magnetic stirring is a classical option in the laboratory for cultures of small volumes. Unlike air bubbling, it is compatible with *in-situ* OD measurement and it has been adopted by the vast majority of the proposed continuous culture systems to which we compare (Table S1B). A specific feature of our system is the precise control of stirring speed (rounds per min), enabling stirring to remain stable over time even if the motors wear out, and to adjust vortex depth so as not to interfere with OD measurement.

The versatility of use and future developments depend largely on electronics, communication, and software architectures. To be easily produced, maintained, and assembled for various continuous culture protocols, the ModuloStat system is designed as a collection of independent devices, complying with Internet of Things principles (^29^). Gervasi *et al*. (^27^) describe the same idea and use similar microcontrollers, but only for peristaltic pumps, not the full system. This architecture is thus a distinctive feature compared to all other systems (Table S1C). The work is divided among boards that are specifically designed to perform subsets of tasks by operating combinations of actuators and sensors (for example temperature management or fluidic transfers). Each board carries a microcontroller that connects to a Wi-Fi network, and an experiment can involve an arbitrary number of boards stacked next to the thermoregulated box (Figure 1). Electric wires connect the boards to the actuators and sensors inside the box for power alimentation and information transfer. After initialization, during an experiment, boards need only a power socket to maintain actuator activity and collect sensor data. On the same Wi-Fi network, possibly in another room, the computer controlling the experiment can interrogate the boards and send new instructions to modify the parameters. Multiple experiments can be run in parallel and controlled from the same computer. New board models can be developed and integrated into the system.

The specifications of the ModuloStat system and their comparison to a set of other previously published systems are summarized Table 1 and Tables S1A-C.

### Digital fabrication of the building blocks for continuous evolution in bioreactors

In line with FabLab’s founding principles of enabling the local production of items conceived elsewhere (^25^), the ModuloStat relies on easily sourced standard components and reproducible digital fabrication (Table 2). The full documentation including program codes, model sources, digital fabrication files, complete list of materials, and complete assembly guide with step-by-step pictures (illustrated Figure S1), is accessible online (https://forge.inrae.fr/modulostat/modulostat).

**Table 2:** Function and composition of modules composing the computer controlled continuous culture modular system.

| Module | Behavior |  |  | Components |  |  |  | Extra informations and typical usage |
| --- | --- | --- | --- | --- | --- | --- | --- | --- |
|  | Passive | Sensor | Actuator | Standard (screws, electronics, tubes, ...) | 3D printed | Laser cut | Custom PCB |  |
| Box | ● |  |  | ● | ● | ● |  |  |
| Bioreactor assembly | ● |  |  | ● | ● | ● |  | withstand autoclave |
| Bioreactor holder | ● |  |  | ● | ● | ● |  |  |
| Magnetic stirring |  | ● | ● | ● | ● |  | ● | 600 to 1000 rpm |
| Air renewal |  |  | ● | ● |  |  |  | through filter |
| Optical density |  | ● |  | ● | ● |  | ● | 0.01 to 1 OD<sub>940</sub> |
| Temperature control |  | ● | ● | ● | ● |  | ● | 37°C |
| Peristaltic pump |  |  | ● | ● | ● | ● |  | up to 100 mL/h |
| Electronic control |  |  |  | ● | ● |  | ● | Based on ESP32 |

Digitally fabricated elements of the ModuloStat range from the caps of the bioreactor vessels to the thermoregulated box, passing by the peristaltic pump heads (Figure 2 and Figure S2). The autoclavable screw cap of the culture vial is made of 3D-printed polypropylene with a laser-cut silicone seal. It is designed to hold four or five stainless-steel tubes for liquid and air transfers (Figure 2A). The peristaltic pump heads are made of 3D-printed PETG and have four standard-size metal rolling bearings that compress the tube and displace the liquid. The thermoregulated box, which holds diverse system components such as liquid and air pumps, magnetic stirrers, and heating elements, is designed as two pieces that assemble into a closed box (Figure 2B). The faces of these two pieces consist of layers cut by a laser and stacked together. Clear poly(methyl methacrylate) (PMMA) provides structural rigidity; cork provides heat and light insulation. The faces are held together using nuts, bolts, and 3D-printed corners made of polyethylene terephthalate glycol (PETG). All of the 3D-printed parts were designed to be printed without support, which limits the amount of manual intervention in the fabrication process. The used printer and laser cutter have the characteristics of standard FabLab equipment.

**Figure 2:**
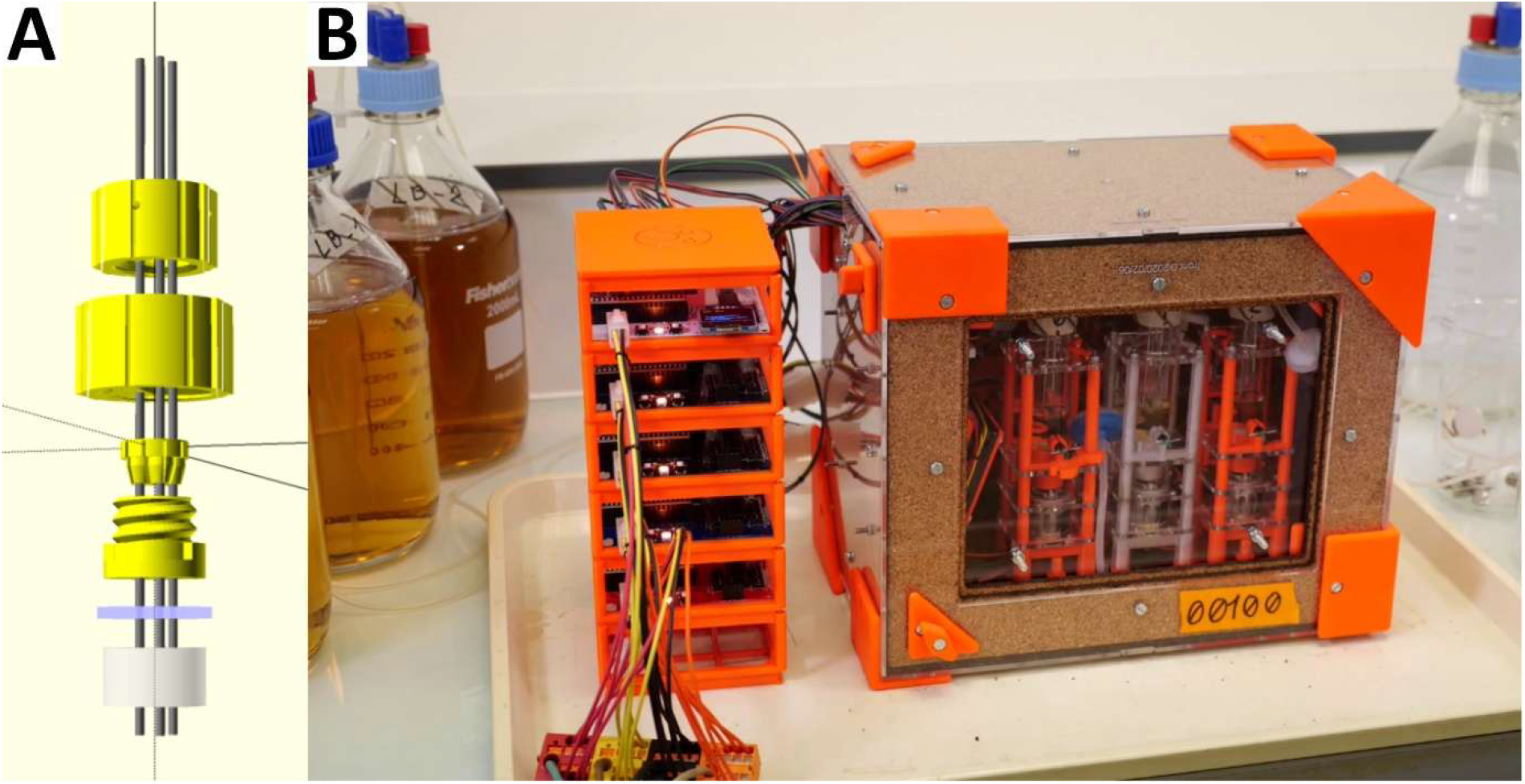
From digital fabrication to continuous culture. **A** OpenScad 3D rendering of the mini-bioreactor screw-cap. Parts to be 3D-printed in polypropylene (yellow) and to be laser-cut in silicone (light blue) maintain the stainless-steel tubes (gray) over the neck of the glass vial (light gray). **B** Global view of the ModuloStat. The thermoregulated box (represented closed, dimensions 30 × 20 × 25 cm) is made from 3D-printed PETG (orange), laser-cut PMMA (clear sheets), and laser-cut cork (light brown textured sheets). The electronic boards (IoT devices) that control the experiment form a stack on the left of the box. Feeding and waste bottles (here 2 L) are at room temperature.

In practice, up to three mini-bioreactor holders, six peristaltic pumps, one air pump, and one heating system can fit inside the thermoregulated box. Additional peristaltic pumps can be placed outside, if needed. The standard vial used as a bioreactor has a volume of 20 mL. This allows for an effective volume between 6 and 16 mL, with sufficient vertical space for *in-situ* optical density measurements between the ferromagnetic bar and the vortex for the smallest volume and enough air volume above the culture for the largest one. The fluidic pumps activate every 30 seconds by default and can in this condition deliver precisely from 1.5 to 25 ml/h by compressing a 1.6 mm inner-diameter tubing (PharMed® BPT). For a 7 mL culture at steady state, the lower flow rate would correspond to steady-state generation times of more than 3h and the higher flow rate is enough to flush any population of planktonic bacteria (see Supplementary Text). This range can be modified by using different tubing diameters or by changing the pump activation frequency. On the same board, the pumps can be activated either independently or sequentially. The thermoregulation system is designed to maintain an internal temperature of 37°C at a typical room temperature of ≈20°C. Since the thermoregulation system only heats, the room temperature would need to be decreased to effectively maintain a temperature close to 20°C. Conversely, to operate at high temperatures, the PETG parts located inside the box can be 3D-printed in a different material, such as polycarbonate (^30^).

### Computer control using IoT principles

The ModuloStat boards contain the electronics needed to provide a basic level of control on a set of functional modules (actuators and/or sensors), allowing to perform a subset of tasks for an experiment. In this work, four types of boards controlling different sets of modules have been used (Figure S3). Each board consists of a printed circuit (PCB) of format 89 × 100 mm, whose design is split into a part common to all types of boards and a part specific to each type of board. The common part hosts power-wire connectors and an ESP32 (Espressif), which is a fast programmable microcontroller with wireless communication capabilities and enough onboard memory to withstand a single C++ program with object representation of all the types of actuators or sensors. Micro-switches allow for on the fly choice of the Wi-Fi network, identification of an experiment, identification of a recipe matching the variable part of the board. Several switches also serve to distinguish identical boards belonging to the same experiment (Figure S3). The recipe determines how the common C++ program should be executed by describing the board components and their initialisation parameters. They are stored as easy-to-modify JavaScript Object Notation (JSON) files. When powered, the board automatically connects to a configured wireless network and starts working. A small plug-and-play screen can be connected to the PCB to access a board-level read-only user interface. Updating board parameters is done through Wi-Fi.

Orchestrating an experiment can involve up to three levels of control over the behavior of the boards (Figure S4). The boards maintain their configured behavior and act as web servers that answer HTTP requests to update parameter values and return measured values. Information is sent via Wi-Fi using the HTTP standard communication protocol with a RESTful API design based on a JSON data transfer format. The computer code on the boards represents the first layer of system control and makes it possible to operate the system manually by HTTP requests. However, to orchestrate the boards’ behavior in more complex experiments, which are typical of continuous cultures, we used programs that run on a Raspberry Pi mini-computer. For simplicity, we also used this mini-computer to deploy the Wi-Fi network. The second layer of control consists of a program that periodically checks the status of the boards, synchronizes their running parameters with those provided in a simple configuration text file, and gathers the latest measured sensor values to compile them into a large table that records the entire experiment history. This second layer of control also restores parameters after a voluntary or involuntary shutdown of the boards and automatically formats and sends requests to change parameters after a manual modification of the configuration text file. This ensures that simple systems, such as chemostats, can run for weeks with manual control and simple graphic generation to display system states over time. A third layer can be added for more advanced control, such as automatically adjusting a parameter of the culture based on the collected OD data. In this case, a program automatically changes the configuration text file. For example, it changes the bioreactor dilution rate when the OD crosses a threshold level. This layered control provides great flexibility in using the ModuloStat’s functional modules. Combining low-level control at the board level with externalization of high-level control on a distant computer scales well and allows for the implementation of sophisticated protocols.

### Experimental validation of functional modules

We performed several experiments to validate and characterize the key elementary properties of the ModuloStat’s functional modules: OD monitoring in the near-infrared, setting of liquid flow rate using peristaltic pumps, setting of the bioreactor dilution rate, temperature management, and magnetic stirring.

The OD of the culture is measured in the near-infrared spectrum at 940 nm rather than in the visible spectrum at the more common wavelength of 600 nm. For serial dilutions, we compared our *in-situ* measurements with those of a bench apparatus used to measure the optical density of microbiological cultures in laboratories (Figure 3A). The results were very similar, confirming the ability of our module to monitor biomass changes in the near-infrared. When compared directly in a multi-wavelength spectrophotometer, the near-infrared measure is twice lower than the visible measure and, possibly for this reason, is slightly more linear in the high biomass range (Figure S5).

**Figure 3:**
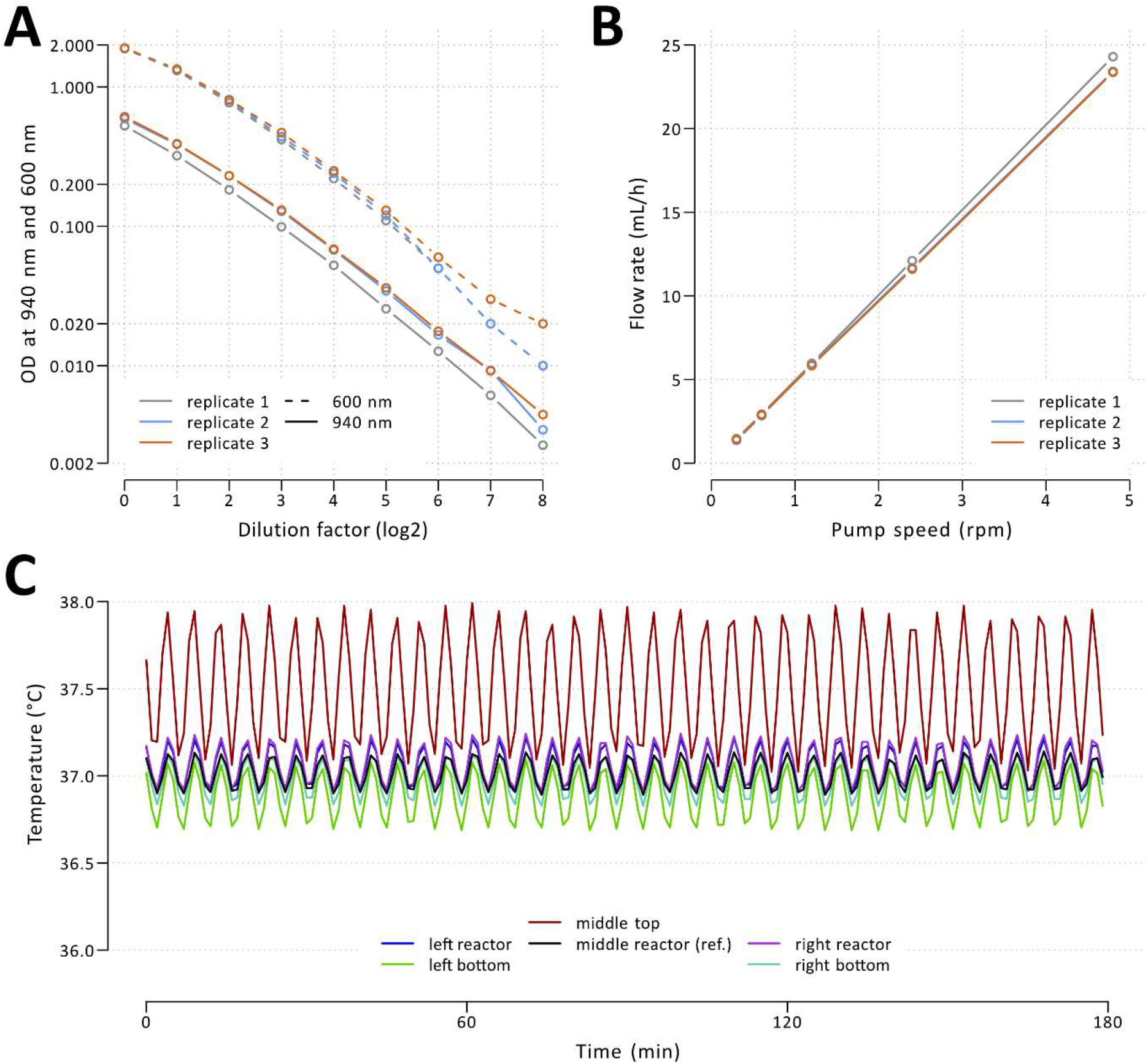
Experimental validation of functional modules. **A** Comparison between *in situ* OD measurements at 940 nm and measurements made at 600 nm with a bench density meter. **B** Relationship between measured flow rate and imposed peristaltic pump speed. **C** Temperature measured at six points inside the box over time, with a target set to 37°C.

The peristaltic pumps, which are composed of a newly designed digitally manufactured pump head and a stepper motor, were evaluated on their ability to deliver a precise liquid flow rate. To accomplish this, we measured the flow rate at various rpm with the module’s default settings, i.e., without altering the rotational speed and only changing the duration of the move executed every 30 seconds. The results revealed high linearity and reproducibility. One rpm corresponded to 4.93 ml/h (Figure 3B). However, we observed that at very low rpm, the flow rate was slightly lower than expected (approximately 5% lower at 0.3 rpm) (Figure S6). Our initial hypothesis was that this could be due to a small backlash movement caused by the elasticity of the tubing when the motor tension shuts down after the move is executed (^27^). However, the same observation was made when the motors were kept electrically blocked between moves. Another possible reason is the gas porosity of the silicone tubing used on both sides of the Pharmed BPT section in the pump head, or the evaporation from the mini-bioreactor due to forced air renewal.

The dilution rate (δ) is the central parameter of a bioreactor regime. It is tied to flow rate (f) and volume (v) by the relationship δ = φ/v (see Supplementary Text). The procedure for setting the volume of the culture consists of adjusting the height of the stainless-steel tubing’s mouth, through which the medium is extracted from the bioreactor and which works as an overflow. The liquid surface tension at the mouth and the vortex caused by stirring make this setting non-trivial (Figure S7). Uncertainty in volume causes uncertainty in dilution rate, which can deviate slightly from its target value and, consequently, affect the growth rate of cells at culture steady state. Since we have already established the linearity of our OD measurement with biomass concentration, we can also directly estimate the dilution rate using conditions of pure dilution, i.e. without growth. This allowed us to check the precision with which the dilution rate could be set. For a target dilution rate of 0.0122 min^-1^, which corresponds to a steady-state generation time of 56.8 min, we measured a mean dilution rate of 0.0117 min^-1^ across three bioreactors (Figure S8). With a coefficient of variation of 2.7% and a relative deviation of S4% with respect to the user-defined target, this result indicates that the current calibration procedure makes it possible to set the dilution rate with reasonable precision.

Changes in temperature over time and differences between bioreactors should be minimized because microbial growth rates vary considerably with temperature. The temperature management board integrates a heating resistor and a high-precision digital temperature sensor to implement a simple on-off controller that raises and maintains the temperature near the user-defined target. We took advantage of our system’s modularity in this test to add a board and measure the temperature at six positions simultaneously Figure 3C). Initial results obtained with a single fan led us to install a second fan to homogenize the temperature inside the box (Figure S9). With this setup and a target of 37°C, we measured that the temperature near the three bioreactors oscillated within a narrow range of 36.7 to 37.3°C with a period of only ≈4.5 minutes. When air renewal is activated, the air inside the bioreactors is pushed out and replaced with air taken from within the box near the bioreactors.

The culture is homogenized by stirring with a magnetic bar whose rotating speed ranges from 600 to over 1,200 rpm. The onboard electronics automatically adjust the motor’s voltage using pulse width modulation to maintain the desired speed, as measured by a Hall effect sensor. When set to 600 rpm, as appropriate for a culture of 7 mL, the effective speed oscillated within a narrow range of standard deviation 14 rpm (Figure S10).

### Engineering of a bacterium with a zero-biofilm phenotype in chemostat

The first test of the ModuloStat involved implementing the chemostat mode of culture (Figure 4). In a single thermoregulated box, we conducted three independent cultures of *B. subtilis* BSB1. After approximately 6 hours at a dilution rate corresponding to a steady-state generation time of 60 minutes, the biomass, as measured by OD, stopped increasing and even decreased slightly, indicating that the growth rate temporarily decreased slightly below the dilution rate. After approximately 12 hours, we manually increased the dilution rate, which led to a quick decrease of the OD, followed by stabilization. An equilibrium was reached at this point, and the population was continuously propagated under a constant generation time of approximately 30 minutes. However, after another 48 hours, i.e. 96 generations under the imposed dilution rate, the OD signals increased and varied erratically, indicating a loss of equilibrium (Figure 4A).

**Figure 4:**
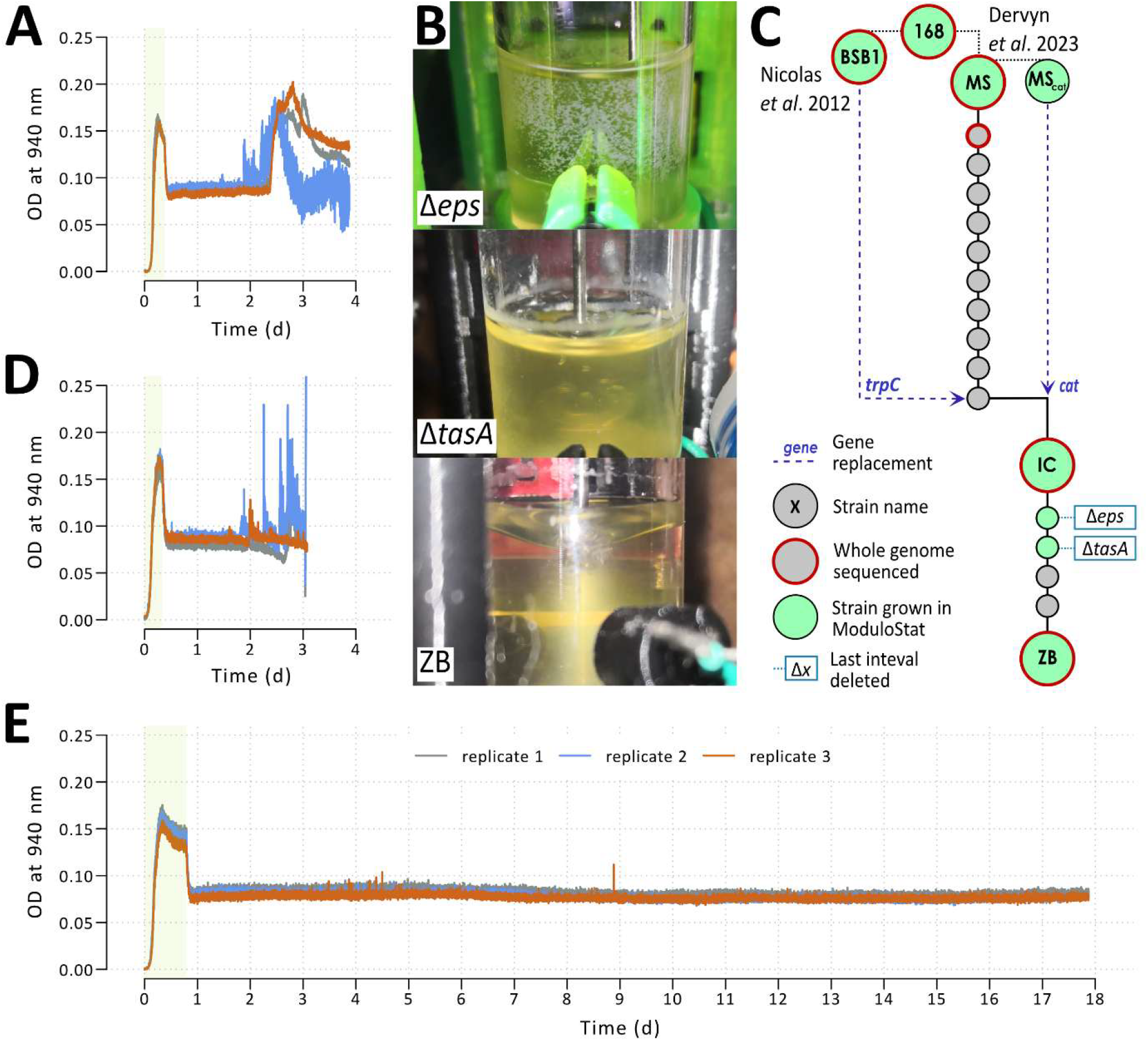
Biofilm formation in chemostat cultures and engineering of a zero-biofilm strain. **A** OD trajectory of chemostat cultures of *B. subtilis* BSB1 strain with a dilution rate corresponding to a steady-state generation time of 30 min. Biofilm formation was detected after two days of culture by anomalies in OD measures. The dilution rate was two times lower during the period indicated by the light green vertical area than during the rest of the culture. OD values were sampled every minute. **B** Images of chemostat cultures with and without biofilm formation: a submerged biofilm formed by the *Δeps* intermediate strain, an emerged biofilm formed by the *ΔtasA* intermediate strain, a culture without biofilm formation (ZB strain). **C** Construction lineage of the zero-biofilm (ZB) strain. **D** The intermediate chassis (IC) strain also formed biofilm after two days of culture. **E** The ZB strain did not form biofilms until run-out of the feeding bottle after 18 days.

Visual examination of the cultures revealed the presence of biofilms on the glass walls of the three mini-bioreactors. These biofilms were of the type of those presented Figure 4B. Like its parental *B. subtilis* 168 strain, the BSB1 strain is a laboratory strain which is not considered a good biofilm former compared to non-domesticated strains such as NCIB 3610 WT or NDmed (^31^). However, the number of generations that have elapsed in the bioreactor has likely enabled the selection of mutants that can evade dilution by adhering to glass walls. This observation aligns with those made in other organisms, such as *E. coli*, which cannot be cultivated in a simple chemostat for extended periods.

The status of *B. subtilis* as a model organism comes with high genetic tractability and good knowledge of gene functions. Therefore, we reasoned that engineering a strain particularly well-suited for long-term continuous evolution by deleting selected groups of genes would be feasible and interesting (Figure 4C, genotypes in Table S2, coordinates of deletion intervals in Table S3). We began this process by removing a preliminary set of genes starting from a master strain (MS), a derivative of BSB1 in which genomic islands that could lead to lysis or genetic instability were already removed (^32^). Using synthetic biology tools we additionally removed genes essential for forming highly resistant spores, which could complicate sterilization of the system; for operating flagella, which could allow the bacterium to travel actively (against the liquid flow) in the flexible tubes connected to the bioreactor; and for producing antibacterials, which could allow antagonistic interactions between cells. After a similar amount of time to that of the BSB1 strain, chemostat cultures inoculated with the resulting strain (IC) started to form biofilms and to exhibit OD instability (Figure 4D). However, the instability patterns may differ between the two strains, as suggested by comparing with Figure 4A.

The genome contains a large region of 15.7 kb that encodes *eps* genes, which play a key role in synthesizing the exopolysaccharides that compose the extracellular matrix of *B. subtilis* biofilms. We removed this region, as well as the *tasA* gene, which encodes a protein that assembles into long fibrils important for biofilm structure and integrity. However, this was not enough to prevent the formation of surface-adhering colonies that could escape the outflow (Figure 4B). Based on sequencing data for clones isolated from various biofilms or aggregates after evolution in a chemostat (not shown) and a scan of the SubtiWiki database (^33^) to identify genes whose functions have been described in the literature as associated with biofilm formation or surface adhesion, we removed three additional *loci*. The first two encode proteins involved in hydrophobic properties (*bslA*) and in the synthesis of poly-gamma-glutamate, a homopolymer that can be part of extracellular matrices (*pgsBC*). The third locus consists of an operon (*ydaJKLMN*), which is lesser known than the eps genes but whose products also mediate the synthesis of exopolysaccharides. The accumulation of deletions did not substantially impact the generation time (Figure S11). When inoculated with this strain, a stable OD was observed in the chemostat until the end of the feeding bottle (Figure 4E), i.e., approximately 800 generations (18 days). We named this strain *B. subtilis* ZB.

### Implementation of various types of continuous cultures with the ModuloStat

To further illustrate the use of the ModuloStat, we implemented more complex continuous culture modes (Figure 5). After the chemostat (Figure 5A), we first considered evolving a resistance to a stressful condition and selected exposure to ethanol, which has multiple effects. Ethanol notably increases membrane fluidity, resulting in uncontrolled transport of solutes, and it binds to some proteins, including enzymes and transcription factors, altering their functions. Alcohol stress is particularly relevant since it occurs in important fermentation conditions, such as biofuel production. Resistance to ethanol has already been studied through experimental evolution in several bacteria using discontinuous cultures, particularly *E. coli* (^34^).

**Figure 5:**
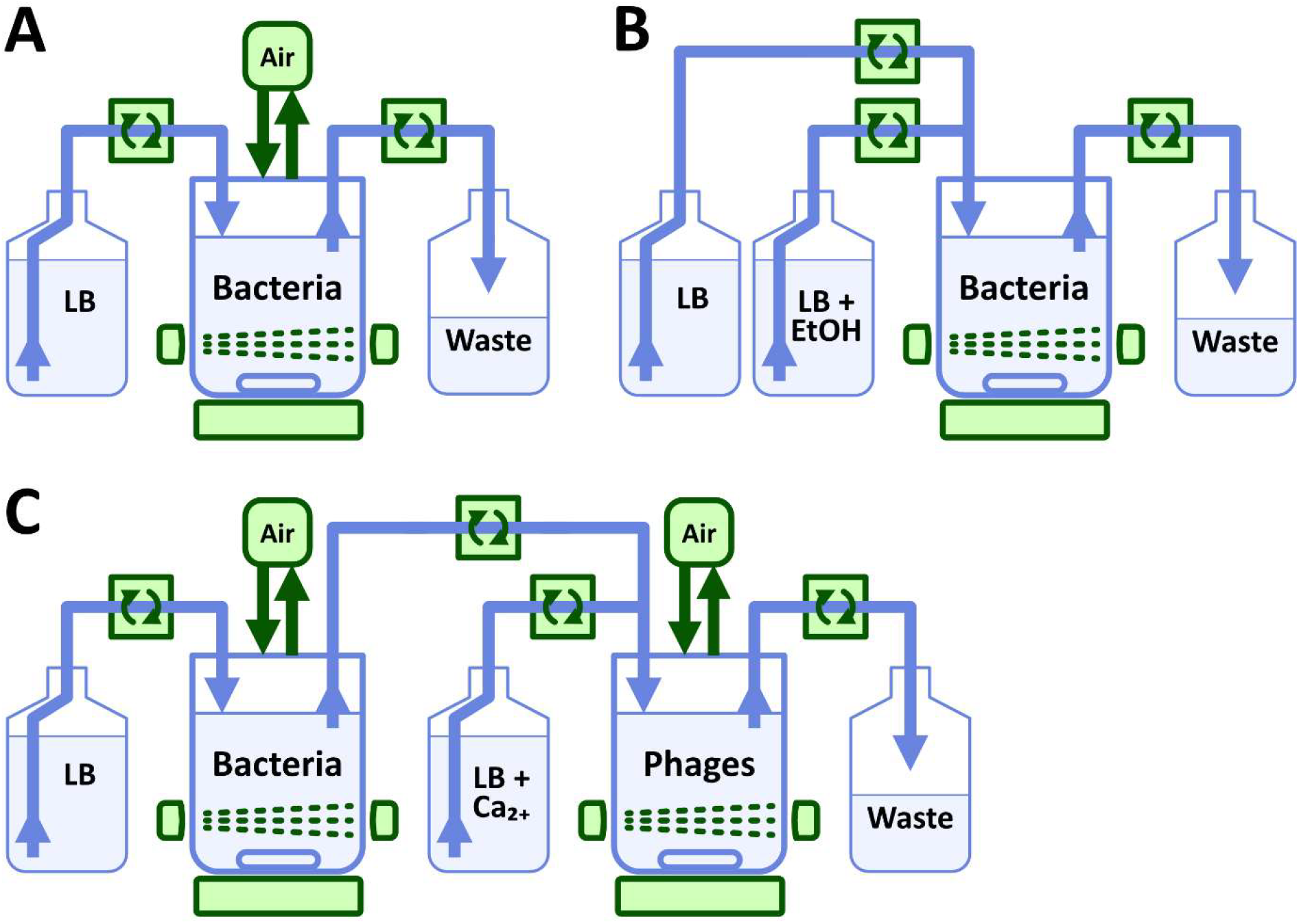
Mini-bioreactor assemblies to exemplify the use of the ModuloStat. **A** Assembly for chemostat and turbidostat cultures. Air pumps were turned off for the chemostat cultures presented in the text. **B** Assembly for the evolution of resistance to ethanol stress. **C** Cascade of two mini-bioreactors for the continuous culture of phages.

Inspired by the morbidostat (^14^), our culture protocol consisted of a constant dilution rate and an increasing ethanol concentration. We conducted the experiment without aerating the bioreactors because preliminary tests indicated that aeration could reduce ethanol toxicity by facilitating its degradation through aerobic metabolism or by evaporation. Each bioreactor was fed by a mixture from two bottles of growth medium: one without ethanol and one in which some of the water was replaced with ethanol (Figure 5B). The proportions of the mixture were automatically adjusted when the OD crossed certain threshold values (Figure 6A). We defined a step of 0.48% for this adjustment in the total ethanol concentration. If the OD was above an upper threshold, the ethanol concentration increased by one step. To prevent extinction or severe bottlenecks, we also defined a lower OD threshold. If the OD was below the lower threshold, the ethanol concentration decreased by one step. We set the dilution rate to achieve a steady-state generation time of 60 minutes. The upper OD threshold corresponded to 75% of the equilibrium OD for a dilution rate twice as high. This means that when the OD stabilizes below this threshold, ethanol has an effect that results in more than a doubling of the generation time. This can be considered mild inhibition of growth.

**Figure 6:**
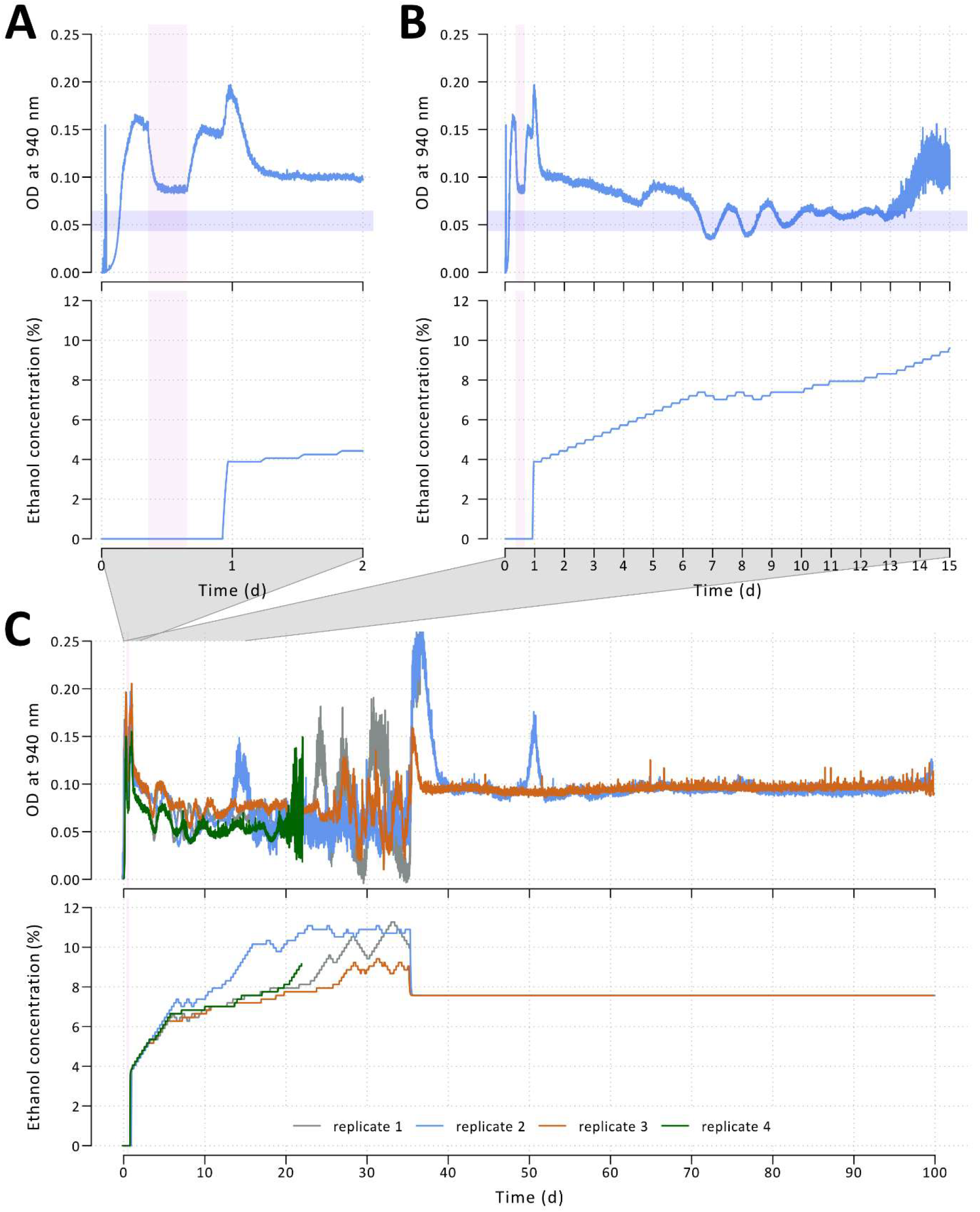
Trajectories of cultures during the evolution of resistance to ethanol stress. **A** OD and ethanol concentration during the first 2 days of cultures of the *B. subtilis* ZB strain, illustrated for one of the four replicates. For this representation, the theoretical ethanol concentration was computed by integrating the proportion of the mixture of the two feeding bottles (with and without ethanol), taking into account the dilution rate in the bioreactor. The dilution rate corresponded to a steady-state generation time of 60 min, except during a brief period with a dilution rate two times higher (light pink vertical area). The upper and lower OD thresholds (75% and 50% of the OD at steady-state generation time of 30 min) used to adjust the concentration of ethanol were determined during this brief period and delimit the light blue horizontal area. A minimum time interval of 6 h is imposed between successive adjustments to the ethanol concentration in the same direction. After the initial ethanol addition, the OD transiently increased before rapidly decreasing. This is likely due to the consumption of remaining oxygen by ethanol before the effects of ethanol stress are seen. **B** same as **A**, but for the first 15 days of the experiment. The OD decreased progressively following successive increases in ethanol concentration until day 4, when an abrupt increase in the OD likely corresponded to a first visible adaptation of the bacteria. On day 6, the OD decreased below the lower threshold, and the ethanol concentration was automatically decreased. On day 7, the OD surpassed the upper threshold, and the ethanol concentration increased automatically. On day 13, the “flake” phenotype appeared in the culture, resulting in a noisier OD signal. **C** OD and ethanol concentration for the four replicates throughout the entire experiment. The “flake” phenotype appeared in each replicate when the ethanol concentration reached approximately 8%. On day 35, the ethanol concentration was fixed at 7.5%, and the culture continued for two replicates whose phenotypes reverted until the input medium ran out on day 100. Represented OD values were sampled every minute for **A** and **B**, every 10 minutes for **C**.

Four independent cultures were inoculated with strain ZB and conducted in parallel in two thermoregulated boxes. The initial ethanol concentration was set to 3.88%, and the automatic adjustment procedure started to gradually increase the concentration (Figure 6A) until the OD stabilized between the upper and lower thresholds (Figure 6B and S12). However, after 14 days or more, when the ethanol concentration reached approximately 8%, we observed an increasingly noisy OD signal (Figure 6B and 6C). Visual inspection of the bioreactors revealed the presence of “flakes” that explained this change (Figure S13). This phenotype made controlling the OD difficult between days 14 and 35. By this time, the ethanol concentration had reached 9.4% to 11.3%, depending on the bioreactor. We decided to halt the automatic adjustment procedure and continue with a fixed ethanol concentration of 7.5%. In two out of four bioreactors, the drop in ethanol concentration reversed the “flake” phenotype, as evidenced by the smoother OD signal. We continued these cultures until day 100. Given the dilution rate, this corresponds to more than 2,000 generations. This experiment, which was conducted without external contamination or antibiotic treatment, confirmed the robustness of the ModuloStat system.

In the case of evolving ethanol resistance, the OD was controlled by adjusting the composition of the growth medium. In contrast, the traditional turbidostat involves controlling the OD by adjusting the dilution rate. To demonstrate the ModuloStat’s versatility, we implemented a culture in which the OD was also maintained below 75% of the equilibrium OD at a generation time of 30 minutes (same thresholds as for the ethanol stress experiment), but by switching between phases of low and high dilution rates. These phases corresponded to theoretical steady-state generation times of 30 and 10 minutes, respectively. Because the bacteria could not achieve a generation time of 10 minutes, the high dilution rate reduced the population mechanically when the threshold was crossed. This occurred approximately every 30 minutes (see Figure S14).

To provide one final example of how the ModuloStat can be used, we assembled a two-bioreactor cascade for the continuous culture of a bacteriophage. Inspired by the Phage-Assisted Continuous Evolution protocol in *E. coli* with its well-known chronic phage M13 (^6^), the assembly consisted of a first bioreactor that produces bacteria to feed the phages in a second bioreactor (Figure 5C). The first bioreactor was run as a chemostat, and the second was referred to as the “lagoon”. In this setup, we cultivated *B. subtilis* ZB and the lytic phage SPP1 (^35^). The two bioreactors were placed in the same thermoregulated box, and the lagoon was also fed by a second bottle of medium containing a high concentration of calcium. Calcium is required for infection by SSP1, and its absence in the first bioreactor ensures that the phage cannot contaminate the upstream culture. The second bottle also allows for a higher dilution rate in the lagoon than in the chemostat. In practice, the dilution rate of the chemostat is set such that the bacterial generation time at steady state is approximately 30 minutes. The outflow of the chemostat constitutes 50% of the lagoon’s feeding, resulting in a dilution rate that is twice higher in the lagoon than in the upstream chemostat. This ensures that a resident population of bacteria cannot establish in the lagoon. Consequently, lineages of phage-resistant bacteria, which emerge as a result of mutations, should be continuously eliminated.

After reaching equilibrium with a generation time of 30 minutes, the OD in the upstream chemostat remained stable until the feeding bottle ended on day 19. Prior to phage inoculation, the ODs of the upstream and downstream bioreactors were similar (Figure 7). Phage inoculation on day 2 lowered the optical density in the lagoon due to bacterial lysis. An apparent equilibrium ensued and persisted until the end of day 6. On day 7, the OD in the lagoon began oscillating with a period of approximately 5 hours. These oscillations were somewhat unexpected, yet they appeared repeatedly when testing different dilution parameters (not shown). The delayed appearance of the oscillations may suggest that they result from an evolution of the phage population.

**Figure 7:**
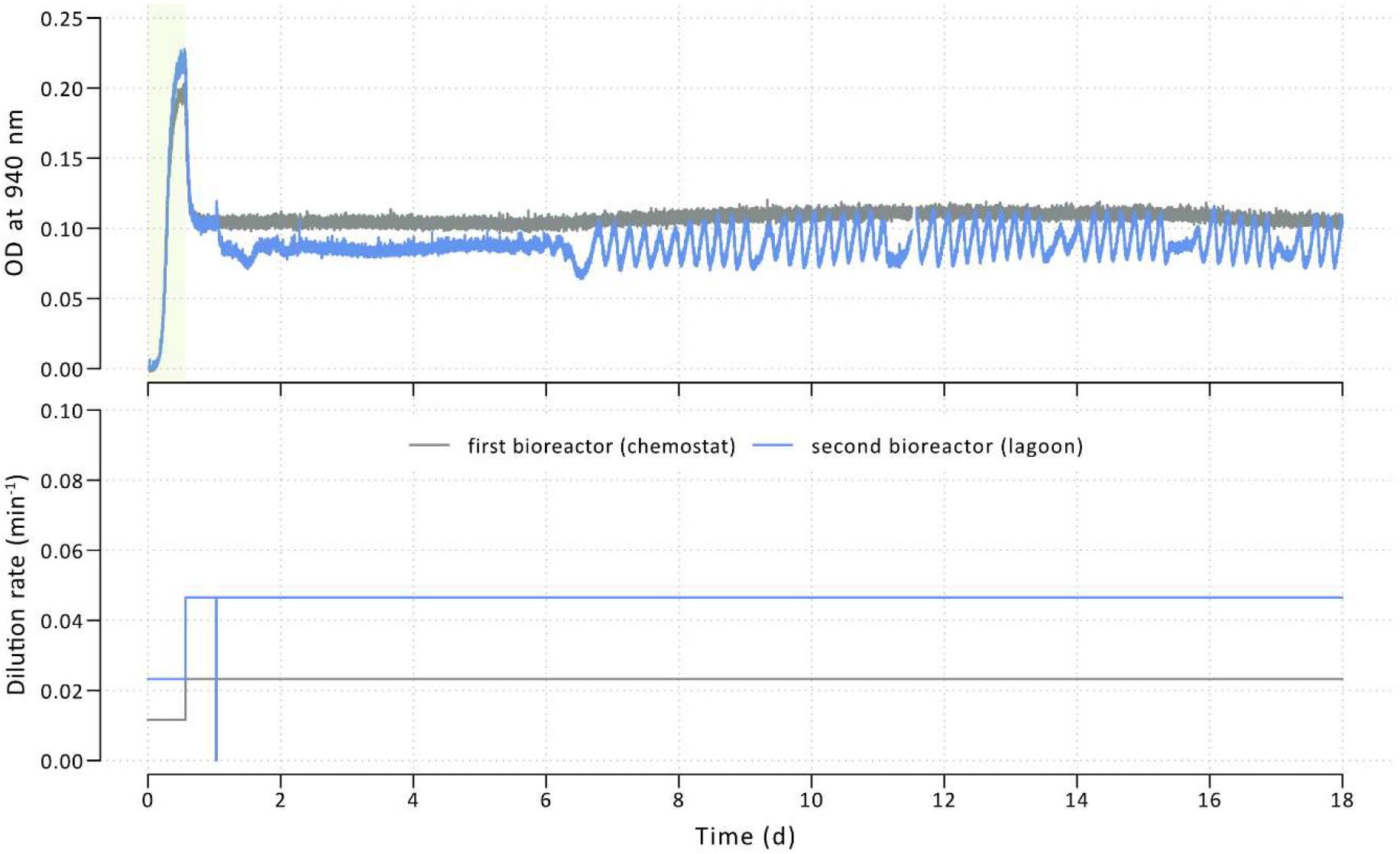
Continuous culture of SPP1 phage using a cascade of two mini-bioreactors. OD trajectories and dilution rates are represented for the two bioreactors. In the first bioreactor, the *B. subtilis* ZB strain was cultivated in a chemostat mode. In the second bioreactor (“lagoon”), SPP1 phages were inoculated after one day of culture. The dilution rate of the chemostat corresponded to a steady-state generation time of 30 min, except for the period indicated by the light green vertical area, when it was two times lower. Represented OD values were sampled every minute.

### Conclusion and perspectives

The ModuloStat is a modular, open-source, framework that facilitates continuous culture in mini-bioreactors. The ModuloStat system is based on digital fabrication tools and programmable electronics that are easily accessible. Maintaining a culture is divided into tasks assigned to dedicated printed circuit boards. According to Internet of Things principles, each board autonomously operates a set of sensors and actuators and can receive and send information. The boards are stacked to implement complex behaviors.

The modularity of the ModuloStat stems from its architecture. A first level of modularity lies in the freedom to combine an arbitrary number of the four types of boards described here. A second level of modularity comes for the possibility of designing new boards that reassociate already developed functional modules. Using such new boards only requires describing a new recipe in a JSON text file. A third level of modularity stems from the design of the boards and of the microcontroller code, which are intended to easily accommodate new functional modules. The corresponding C++ object in the microcontroller code would then be programmed using the existing modules as models. The new boards will be integrated with others in a common fleet of IoT objects.

The *B. subtilis* ZB strain developed in this work is to our knowledge the first bacterium designed to have a zero-biofilm phenotype when grown in continuous culture. *B. subtilis* ZB could then become a model for long-term evolution in bioreactors and a chassis for synthetic biology applications based on adaptive laboratory evolution. Transferring in this strain the genetic systems providing conditional hypermutator phenotypes in *B. subtilis* (^36^) may further allow to accelerate and better control evolution in the laboratory.

The ModuloStat’s versatility and robustness have been illustrated through the implementation of various continuous culture modes implying different mini-bioreactor fluidic topologies. New culture modes can be conceived with the existing functional modules. Digital fabrication and parametric 3D modeling allow users to readily customise shapes and dimensions to meet new technical specifications, such as a different bioreactor vessel size or a different type of motor. The new functional modules that may be conceived in the future, could include *in-situ* fluorescence sensors to monitor the expression of reporter genes and actuators that operate syringe pumps or pinch valves to extend the fluidic design possibilities.

## METHODS

### ModuloStat construction

Assembly instructions, bill of materials, digital fabrication files (for 3D printing, laser cutting and circuit boards) along with programs and 3D model sources are available at https://forge.inrae.fr/modulostat. 3D parametric models were designed using OpenScad software and fabricated using a Prusa i3 MK3S+ 3D printer and a FabCreator FabKit Mk 6 laser cutter. Circuit boards were designed using Fritzing and ordered to a commercial manufacturer (JLCPCB).

### Functional modules characterisation

A ModuloStat system was configured for three parallel chemostats fed with LB (1.04 rpm) with active temperature control set to 37°C, two active fans, magnetic stirring at 600 rpm, and forced air renewal turned on. Those parameters were set for the entire characterisation unless stated otherwise.

The peristaltic pumps were tested for the transfer of LB medium at speeds ranging from 0.3 to 2.4 rpm. These speeds were achieved by executing a number of steps between 30 and 480 every 30 seconds. The output flow was collected in 15 mL Falcon tubes. After reaching 28,800 steps, which represented 144 complete rotations of a peristaltic pump head (200 steps per turn), the collected liquid was weighed on a precision scale. For each replicate, the coefficient linking the measured flow rate (in ml/h) to the speed of the pump (in rpm) was estimated using the R function “lm” without intercept. The conversion factor from rpm to ml/h was calculated as the mean of the coefficients from the three replicates.

To characterize OD measurements, saturated cultures of *B. subtilis* grown in LB were mixed with 1mg/mL chloramphenicol to stop cell division. Non-growing cultures were then pipetted in the emptied mini-bioreactors with feeding and outflow pumps turned off. OD measures (940 nm) were taken in the closed box when target temperature was reached. Half of the volume of each bioreactor was then collected to be measured with an Ultrospec 10 Amersham Biosciences density meter (600 nm) and an equal amount of the LB medium supplemented with chloramphenicol was added resulting in a two-fold dilution. These steps were repeated eight times until a 256-fold dilution. OD blank references were taken on LB medium supplemented with chloramphenicol.

For temperature characterisation, an additional board with four extra ambient temperature probes was used to measure the temperature variation throughout the box.

### Construction of the *B. subtilis* zero-biofilm (ZB) strain

All strains used in this work derived from *B. subtilis* 168 (Figure 4C, Table S2). *B. subtilis* BSB1 is a prototroph version of *B. subtilis* 168 (^37^) and MS is a tryptophan auxotroph genome-reduced strain (^32^).

The *B. subtilis* IC and ZB prototroph strains were obtained by iterative accumulation of deletions starting from MS using a method that we previously developed (^32^). Briefly, it consisted in transforming *B. subtilis* (^38^) to introduce a deletion by homologous replacement (pop-in) of a targeted chromosome region by a deletion cassette (*upp ble* λC1 or *upp spc* λC1); the phleomycin or spectinomycin resistance gene in the cassette serving for selection of integration. Cassette eviction (pop-out) occurred by spontaneous recombination between two directly-repeated sequences included in the design and selected by restoring either a neomycin or a chloramphenicol resistance phenotypes (resp. in strains carrying λPr-neo and λPr-cat). The strain obtained after the pop-out was used for the next deletion step resulting in the strain collection described in table. When required, antibiotics phleomycin, neomycin, chloramphenicol and spectinomycin were added at final concentrations of 8, 15, 5 and 100 μg/mL, respectively. Strains BSB1, MS, IC and ZB were genome sequenced (Eurofins).

### Sterilization, optical blank, and inoculation in the ModuloStat

Mini-bioreactors configured at 7 mL of culture volume were connected to input media filled with fresh LB medium and waste containers, autoclaved 20 min at 120°C (or 30 min at 125°C when 10 L bottles were used), and installed in the thermoregulated box.

Automatic experiment management was launched with the desired culture parameters. The optical density blank is measured *in-situ* through LB media when the set temperature is reached.

*B. subtilis* strains were streaked on plate from frozen stocks (−70°C), and isolated colonies served to inoculate overnight precultures of 10 mL LB medium at 37°C under constant shaking. Precultures were then diluted 100 times in 10 mL LB medium for 4 hours. A 35 µL volume from these second precultures served to inoculate the bioreactor. Inoculation was done in sterile conditions under a microbiological safety cabinet with the ModuloStat boards temporarily electrically disconnected.

### Continuous culture protocols

The various mini-bioreactor assemblies used in the work and the corresponding on-off status of the air pumps are represented in Figure 5. The target temperature was set to 37°C and the magnetic stirring to 600 rpm. The dilution rates are specified in the legends of the figures representing the OD trajectories.

For chemostat and turbidostat cultures (Figure 5A), each mini-bioreactor was fed from a 5 L bottle of LB.

For the evolution of ethanol resistance (Figure 5B), each mini-bioreactor fed from a bottle of 10 L of LB medium and one bottle of 8 L of concentrated (1.25x) LB medium. After autoclaving, the concentrated LB medium was complemented by 2 L of 96% ethanol resulting in a bottle of 10 L of LB (1x) at 19.2% ethanol. The OD thresholds used to adjust the concentration of ethanol were determined independently for each mini-bioreactors.

For the continuous culture of phages (Figure 5C), the fluidic circuit consisted of a cascade of two bioreactors fed from a bottle of 5 L of LB for the first bioreactor (chemostat) and also from a bottle 5 L of LB with 10 mM CaCl_2_. This second feeding bottle served as a complementary medium input for the second bioreactor containing the phages (the “lagoon”). The effluent from the first chemostat and the LB with CaCl_2_ were transferred to the second chemostat with the same flow rates. Phages were inoculated in the second bioreactor with a multiplicity of infection of 100.

## SUPPORTING INFORMATION

The supporting information consists of a single pdf (ModuloStat_Supplementary.pdf) document containing:

- a short Supplementary Text introducing the basic mathematical equations on microbial growth in bioreactors useful to understand the main text;
- the Supplementary Tables S1-S3 cited in the main text;
- the Supplementary Figures S1-S14 cited in the main text.

## ACKNOWLEDGEMENTS

We would like to thank Paulo Tavares for the kind gift of SPP1 phage and guidance while working with it. We would also like to thank Romain Di Vozzo for introducing us to digital fabrication and for facilitating our access to the UPSaclay FabLab (www.digiscope.fr/en/platforms/fablab). We are also grateful to the Pépinière Numérique at INRAE for granting us access to their digital fabrication resources. Additionally, we would like to thank the INRAE MIGALE bioinformatics facility (https://doi.org/10.15454/1.5572390655343293E12) for the computational resources they provided.

## CONTRIBUTIONS

C.G., E.D., M.J. and P.N. conceived the project and designed the experimental plan. C.G. and P.N. conceived and produced the ModuloStat system. C.G. wrote the system documentation. E.D. and M.C. engineered the zero-biofilm strain. C.G, E.D., J.V., I.T., E.B. and P.N. performed the experiments. S.D. worked on the graphical representation of the data. C.G., E.D., M.J. and P.N. interpreted the results and wrote the manuscript with the contribution of all authors.

## DATA AVAILABILITY

Websites and source code freely accessible at https://modulostat.maiage.inrae.fr/ or https://doi.org/10.57745/BPIUT7.

## REFERENCES

(1) Monod, J. Annales de l’Institut Pasteur : Journal de Microbiologie / Publiées Sous Le Patronage de M. Pasteur Par E. Duclaux | 1950-07-01 | Gallica. 1950, 390–410.

(2) Novick, A.; Szilard, L. Description of the Chemostat. Science 1950, 112 (2920), 715–716. 10.1126/science.112.2920.715.

(3) Kjeldgaard, N. O.; Maaloe, O.; Schaechter, M. The Transition between Different Physiological States during Balanced Growth of Salmonella Typhimurium. J. Gen. Microbiol. 1958, 19 (3), 607–616. 10.1099/00221287-19-3-607.

(4) Narayanan, C. M.; Narayan, V. Biological Wastewater Treatment and Bioreactor Design: A Review. Sustain. Environ. Res. 2019, 29 (1), 33. 10.1186/s42834-019-0036-1.

(5) Mitra, S.; Murthy, G. S. Bioreactor Control Systems in the Biopharmaceutical Industry: A Critical Perspective. Syst. Microbiol. Biomanufacturing 2022, 2 (1), 91–112. 10.1007/s43393-021-00048-6.

(6) Esvelt, K. M.; Carlson, J. C.; Liu, D. R. A System for the Continuous Directed Evolution of Biomolecules. Nature 2011, 472 (7344), 499–503. 10.1038/nature09929.

(7) Gresham, D.; Dunham, M. J. The Enduring Utility of Continuous Culturing in Experimental Evolution. Genomics 2014, 104 (6 0 0), 399–405. 10.1016/j.ygeno.2014.09.015.

(8) Toprak, E.; Veres, A.; Yildiz, S.; Pedraza, J. M.; Chait, R.; Paulsson, J.; Kishony, R. Building a Morbidostat: An Automated Continuous-Culture Device for Studying Bacterial Drug Resistance under Dynamically Sustained Drug Inhibition. Nat. Protoc. 2013, 8 (3), 555–567. 10.1038/nprot.nprot.2013.021.

(9) Smith, H. L.; Waltman, P. The Theory of the Chemostat: Dynamics of Microbial Competition. Cambridge University Press, 1995.

(10) Li, Z.; Liu, B.; Li, S. H.-J.; King, C. G.; Gitai, Z.; Wingreen, N. S. Modeling Microbial Metabolic Trade-Offs in a Chemostat. PLOS Comput. Biol. 2020, 16 (8), e1008156. 10.1371/journal.pcbi.1008156.

(11) Wills, C.; Phelps, J. Functional Mutants of Yeast Alcohol Dehydrogenase Affecting Kinetics, Cellular Redox Balance, and Electrophoretic Mobility. Biochem. Genet. 1978, 16 (5), 415–432. 10.1007/BF00484208.

(12) Lovitt, R. W.; Wimpenny, J. W. T. The Gradostat: A Bidirectional Compound Chemostat and Its Application in Microbiological Research. Microbiology 1981, 127 (2), 261–268. 10.1099/00221287-127-2-261.

(13) Mutzel, R.; Marliere, P. Method and Device for Selecting Accelerated Proliferation of Living Cells in Suspension. Patent US-6686194-B1, February 3, 2004.

(14) Toprak, E.; Veres, A.; Michel, J.-B.; Chait, R.; Hartl, D. L.; Kishony, R. Evolutionary Paths to Antibiotic Resistance under Dynamically Sustained Drug Selection. Nat. Genet. 2012, 44 (1), 101–105. 10.1038/ng.1034.

(15) Leyn, S. A.; Zlamal, J. E.; Kurnasov, O. V.; Li, X.; Elane, M.; Myjak, L.; Godzik, M.; de Crecy, A.; Garcia-Alcalde, F.; Ebeling, M.; Osterman, A. L. Experimental Evolution in Morbidostat Reveals Converging Genomic Trajectories on the Path to Triclosan Resistance. Microb. Genomics 2021, 7 (5), 000553. 10.1099/mgen.0.000553.

(16) Dößelmann, B.; Willmann, M.; Steglich, M.; Bunk, B.; Nübel, U.; Peter, S.; Neher, R. A. Rapid and Consistent Evolution of Colistin Resistance in Extensively Drug-Resistant Pseudomonas Aeruginosa during Morbidostat Culture. Antimicrob. Agents Chemother. 2017, 61 (9), 10.1128/aac.00043-17. 10.1128/aac.00043-17.

(17) Wong, B. G.; Mancuso, C. P.; Kiriakov, S.; Bashor, C. J.; Khalil, A. S. Precise, Automated Control of Conditions for High-Throughput Growth of Yeast and Bacteria with eVOLVER. Nat. Biotechnol. 2018, 36 (7), 614–623. 10.1038/nbt.4151.

(18) Marlière, P.; Patrouix, J.; Döring, V.; Herdewijn, P.; Tricot, S.; Cruveiller, S.; Bouzon, M.; Mutzel, R. Chemical Evolution of a Bacterium’s Genome. Angew. Chem. Int. Ed. 2011, 50 (31), 7109–7114. 10.1002/anie.201100535.

(19) Miller, A. W.; Befort, C.; Kerr, E. O.; Dunham, M. J. Design and Use of Multiplexed Chemostat Arrays. J. Vis. Exp. JoVE 2013, No. 72, e50262. 10.3791/50262.

(20) Takahashi, C. N.; Miller, A. W.; Ekness, F.; Dunham, M. J.; Klavins, E. A Low Cost, Customizable Turbidostat for Use in Synthetic Circuit Characterization. ACS Synth. Biol. 2015, 4 (1), 32–38. 10.1021/sb500165g.

(21) Guarino, A.; Shannon, B.; Marucci, L.; Grierson, C.; Savery, N.; di Bernardo, M. A Low-Cost, Open-Source Turbidostat Design for in-Vivo Control Experiments in Synthetic Biology. IFAC-Pap. 2019, 52 (26), 244–248. 10.1016/j.ifacol.2019.12.265.

(22) McGeachy, A. M.; Meacham, Z. A.; Ingolia, N. T. An Accessible Continuous-Culture Turbidostat for Pooled Analysis of Complex Libraries. ACS Synth. Biol. 2019, 8 (4), 844–856. 10.1021/acssynbio.8b00529.

(23) Steel, H.; Habgood, R.; Kelly, C. L.; Papachristodoulou, A. In Situ Characterisation and Manipulation of Biological Systems with Chi.Bio. PLOS Biol. 2020, 18 (7), e3000794. 10.1371/journal.pbio.3000794.

(24) Gopalakrishnan, V.; Crozier, D.; Card, K. J.; Chick, L. D.; Krishnan, N. P.; McClure, E.; Pelesko, J.; Williamson, D. F.; Nichol, D.; Mandal, S.; Bonomo, R. A.; Scott, J. G. A Low-Cost, Open-Source Evolutionary Bioreactor and Its Educational Use. eLife 2022, 11, e83067. 10.7554/eLife.83067.

(25) Gershenfeld, N. A.; Gershenfeld, A.; Cutcher-Gershenfeld, J. Designing Reality: How to Survive and Thrive in the Third Digital Revolution; Basic Books, 2017.

(26) Diaz, J.; Tomàs, M.; Lefebvre, S. Are Public Makerspaces a Means to Empowering Citizens? The Case of Ateneus de Fabricació in Barcelona. Telemat. Inform. 2021, 59, 101551. 10.1016/j.tele.2020.101551.

(27) Gervasi, A.; Cardol, P.; Meyer, P. E. Open-Hardware Wireless Controller and 3D-Printed Pumps for Efficient Liquid Manipulation. HardwareX 2021, 9, e00199. 10.1016/j.ohx.2021.e00199.

(28) Ekkers, D. M.; Branco dos Santos, F.; Mallon, C. A.; Bruggeman, F.; van Doorn, G. S. The Omnistat: A Flexible Continuous-culture System for Prolonged Experimental Evolution. Methods Ecol. Evol. 2020, 11 (8), 932–942. 10.1111/2041-210X.13403.

(29) Perkel, J. M. The Internet of Things Comes to the Lab. Nature 2017, 542 (7639), 125–126. 10.1038/542125a.

(30) Petersmann, S.; Spoerk, M.; Van De Steene, W.; Üçal, M.; Wiener, J.; Pinter, G.; Arbeiter, F. Mechanical Properties of Polymeric Implant Materials Produced by Extrusion-Based Additive Manufacturing. J. Mech. Behav. Biomed. Mater. 2020, 104, 103611. 10.1016/j.jmbbm.2019.103611.

(31) Dergham, Y.; Le Coq, D.; Bridier, A.; Sanchez-Vizuete, P.; Jbara, H.; Deschamps, J.; Hamze, K.; Yoshida, K.; Noirot-Gros, M.-F.; Briandet, R. Bacillus Subtilis NDmed, a Model Strain for Biofilm Genetic Studies. Biofilm 2023, 6, 100152. 10.1016/j.bioflm.2023.100152.

(32) Dervyn, E.; Planson, A.-G.; Tanaka, K.; Chubukov, V.; Guérin, C.; Derozier, S.; Lecointe, F.; Sauer, U.; Yoshida, K.-I.; Nicolas, P.; Noirot, P.; Jules, M. Greedy Reduction of Bacillus Subtilis Genome Yields Emergent Phenotypes of High Resistance to a DNA Damaging Agent and Low Evolvability. Nucleic Acids Res. 2023, 51 (6), 2974–2992. 10.1093/nar/gkad145.

(33) Elfmann, C.; Dumann, V.; van den Berg, T.; Stülke, J. A New Framework for SubtiWiki, the Database for the Model Organism Bacillus Subtilis. Nucleic Acids Res. 2025, 53 (D1), D864–D870. 10.1093/nar/gkae957.

(34) Swings, T.; Van den Bergh, B.; Wuyts, S.; Oeyen, E.; Voordeckers, K.; Verstrepen, K. J.; Fauvart, M.; Verstraeten, N.; Michiels, J. Adaptive Tuning of Mutation Rates Allows Fast Response to Lethal Stress in Escherichia Coli. eLife 2017, 6, e22939. 10.7554/eLife.22939.

(35) Godinho, L. M.; El Sadek Fadel, M.; Monniot, C.; Jakutyte, L.; Auzat, I.; Labarde, A.; Djacem, K.; Oliveira, L.; Carballido-Lopez, R.; Ayora, S.; Tavares, P. The Revisited Genome of Bacillus Subtilis Bacteriophage SPP1. Viruses 2018, 10 (12), 705. 10.3390/v10120705.

(36) Tanneur, I.; Dervyn, E.; Guérin, C.; Kon Kam King, G.; Jules, M.; Nicolas, P. The Mutational Landscape of Bacillus Subtilis Conditional Hypermutators Shows How Proofreading Skews DNA Polymerase Error Rates. Nucleic Acids Res. 2025, 53 (5), gkaf147. 10.1093/nar/gkaf147.

(37) Nicolas, P.; Mäder, U.; Dervyn, E.; Rochat, T.; Leduc, A.; Pigeonneau, N.; Bidnenko, E.; Marchadier, E.; Hoebeke, M.; Aymerich, S.; Becher, D.; Bisicchia, P.; Botella, E.; Delumeau, O.; Doherty, G.; Denham, E. L.; Fogg, M. J.; Fromion, V.; Goelzer, A.; Hansen, A.; Härtig, E.; Harwood, C. R.; Homuth, G.; Jarmer, H.; Jules, M.; Klipp, E.; Le Chat, L.; Lecointe, F.; Lewis, P.; Liebermeister, W.; March, A.; Mars, R. A. T.; Nannapaneni, P.; Noone, D.; Pohl, S.; Rinn, B.; Rügheimer, F.; Sappa, P. K.; Samson, F.; Schaffer, M.; Schwikowski, B.; Steil, L.; Stülke, J.; Wiegert, T.; Devine, K. M.; Wilkinson, A. J.; Maarten van Dijl, J.; Hecker, M.; Völker, U.; Bessières, P.; Noirot, P. Condition-Dependent Transcriptome Reveals High-Level Regulatory Architecture in Bacillus Subtilis. Science 2012, 335 (6072), 1103–1106. 10.1126/science.1206848.

(38) Anagnostopoulos, C.; Spizizen, J. Requirements for Transformation in Bacillus Subtilis. J. Bacteriol. 1961, 81 (5), 741–746. 10.1128/jb.81.5.741-746.1961.

