## Supporting information (text, tables and figures) for "ModuloStat: An Internet of Things’ Path to Continuous Cultures in Mini-Bioreactors"

**SUPPLEMENTARY DATA**

#### Table des matières

### SUPPLEMENTARY TEXT

**Basic mathematical equations on microbial growth in bioreactors useful to understand the main text**

The text in blue explains the numerical values given in the main text.

***Notations***

growth rate $\mu$ ($h^{-1}$)
generation time, also known as doubling time $T_g$ (h), with $T_g=\log(2)/\mu$
volume of bioreactor v (mL)
flow rate $\phi$ ($mL.h^{-1}$)
dilution rate $\delta$ ($h^{-1}$), with $\delta=\phi/v$
amount of bacteria or biomass per volume unit x (unit can be OD, cfu/mL, …)
amount of growth substrate in the bioreactor s and in the bottle of fresh medium $s_0$

***Relationship between μ and $T_g$***

During the exponential growth phase of a batch culture, the bacterial concentration at time t can be expressed as $x(t)=x_0.\exp(\mu.t)$ or $x(t)=x_0.2^{t/Tg}$.

Since $\exp(\mu.t)=2^{\mu.t/\log(2)}$, the relationship between the growth rate $\mu$ and the generation time $T_g$ is $T_g=\log(2)/\mu$.

For instance, a generation time $T_g = 30$ min corresponds to a growth rate $\mu \approx 0.023$ $min^{-1}$.

***Steady state of the chemostat***

The evolution of the amounts of bacteria x and substrate s in a chemostat can be described by the system of differential equations
$dx/dt = \mu(s).x - \delta.x$
$ds/dt = \delta.s_0 - \gamma.\mu(s).x - \delta.s$ ,
where $\gamma$ is a yield coefficient, describing the number of units of nutrient that are consumed to produce one unit of bacterium.

The growth rate $\mu(s)$ is a decreasing function of the amount of substrate s such as $\mu(0)=0$ and $\mu(s)\geq0$ for s>0. One model is the "Monod function" $\mu(s)=\mu_{max}.s/(K_s+s)$, with $\mu_{max}$ the maximum growth rate (when s $\rightarrow+\infty$) and $K_s$ is the nutrient concentration at which the growth rate is half-maximal.

By definition, the steady state in the bioreactor is characterized by dx/dt = 0 and ds/dt = 0 and thus
$\mu(s).x - \delta.x = 0$
$\delta.(s_0 - s) - \gamma.\mu(s).x = 0$ .

There is a trivial steady state at x=0 and $s=s_0$ and a non-trivial steady state at $s_{eq}$ such as $\mu(s_{eq})=\delta$. The non-trivial steady state is possible only if $\mu(s_0)>\delta$, which occurs when $\mu_{max}>\delta$ and $s_0$ is large enough to sustain growth above $\delta$.

At this non-trivial steady state we have $\delta.(s_0 - s_{eq}) - \gamma.\mu(s_{eq}).x_{eq} = 0$ , and thus $x_{eq}= \delta(s_0 - s_{eq})/\gamma.\mu(s_{eq})$ .

***Relationship between flow rate, dilution rate and culture volume***

The flow rate to maintain the chemostat at dilution rate $\delta$ is $\phi=\delta.v$.

For instance, with a culture volume of 7mL, a dilution rate of 0.023 $min^{-1}$ ($T_g \approx$ 30 min) requires a flow rate of 0.16 $mL.min^{-1}$. A sample of 1 mL can thus be collected from the outflow in 6.25 min.

Conversely, the flow rates between 1.5 and 25 $ml.h^{-1}$ (0.025 $mL.min^{-1}$ and 0.42 $mL.min^{-1}$) measured for the peristaltic pumps correspond to dilution rates of 0.0036 $min^{-1}$ and 0.060 $min^{-1}$, respectively, for a culture volume of 7mL. The lower dilution rate corresponds to a steady state of Tg=194 min (3.24 h). The higher dilution rate is enough to flush any planktonic culture since the Tg would need to be 11.6 min for a population to be maintained.

***Relationship between number of generations and total volume of growth medium consumed at the end of the experiment***

The volume of growth medium consumed to maintain the chemostat at dilution rate $\delta$ during an experimental runtime of length T is $\delta.v.T$.

At steady state, where the generation time is $T_g=\log(2)/\mu$, the runtime needed for n generations is $n.T_g$. Therefore, the volume of fresh medium needed for n generations is $V_n = \delta.v.n.T_g$, which can be rewritten as $V_n = \delta.v.n.(\log(2)/\mu)$ . Since $\mu=\delta$, we have $V_n = v.\log(2).n$.

In other words, the volume needed for one generation is $V_1 = v.\log(2)$, or approximately 0.69v.

For instance, with a 10L bottle of fresh medium, we can perform 10000/(log(2)*7)≈2061 generations in a culture volume of 7mL.

### SUPPLEMENTARY TABLES S1-S3

| Publication | Fluidic [a] | Sterility management |
|---|---|---|
| This work, **ModuloStat** | iPP | Autoclave |
| Gopalakrishnan 2022, **EVE** | iPP | Autoclave |
| Pen 2021 | iPP | Autoclave |
| Gervasi 2021 | iPP, SP | Autoclave |
| Leyn 2021 | iPP, AO | * |
| Steel 2020, **Chio.Bio** | iPP | Autoclave (reactor), 70% ethanol (fluidic), antibiotics (carbenicillin, chloramphenicol, and spectinomycin) |
| Ekkers 2020, **Omnistat** (Ekkers 2022) | iPP | Autoclave |
| McGeachy 2019 | iPP, AO | Autoclave |
| Guarino 2019 | iPP, SP, AO | * |
| Wong 2018, **eVOLVER** (Heins 2019, García-Ruano 2023) | iPP | Autoclave (reactor), bleaching (fluidic) |
| Hoffmann 2017 | iPP, AO | Antibiotics (ampicillin) |
| Dößelmann 2017 | iPP, mPP | Autoclave, antibiotics (experiment on colistin resistance) |
| Takahashi 2015, **Flexostat/Fluorostat** | SP, AO | Autoclave |
| Miller 2013, **Ministat array** | mPP, AO | Autoclave |
| Toprak 2012, **Morbidostat** (Toprak 2013) | iPP | Autoclave |
| Marliere 2011 | * | 5M sodium hydroxide after each culture transfer |
| Tomson 2006, **Adaptastat** | iPP | Antibiotics (rifampicin) |

**Table S1A - Complementary characteristics of already published continuous culture systems.** [a] mPP (multichannel peristaltic pumps with common speed), iPP (individually controlled peristaltic pumps), SP (syringe pumps), AO (air flow driven overflow). * Information not available in publication, supplementary information or associated online data.

| Publication | Organisms [a] | OD wavelength | Stirring | Temperature management [b] |
|---|---|---|---|---|
| This work, **ModuloStat** | *B. subtilis* (b), SPP1 (p) | 940 nm | Magnetic | A (air heater) |
| Gopalakrishnan 2022, **EVE** | *E. coli* | 950 nm | Magnetic | A (external incubator) |
| Pen 2021 | *E. coli* | 590 nm | Magnetic | I (peltier module) |
| Gervasi 2021 | *E. gracilis* | * | Shaking table | * |
| Leyn 2021 | *E. coli* | 654 nm | Magnetic | A (air heater) |
| Steel 2020, **Chio.Bio** | *E. coli* | 650 nm | Magnetic | I (PCB) |
| Ekkers 2020, **Omnistat** (Ekkers 2022) | *L. cremoris* | * | Magnetic | A (water bath) |
| McGeachy 2019 | *S. cerevisiae*, *E. coli* | 940 nm | Magnetic | * |
| Guarino 2019 | *E. coli* | 950 nm | Magnetic | A (external incubator) |
| Wong 2018, **eVOLVER** (Heins 2019, García-Ruano 2023) | *S. cerevisiae*, *S. marcescens* | 900 nm | Magnetic | I (heating block) |
| Hoffmann 2017 | *E. coli* | 650 nm | Magnetic | A (external incubator) |
| Dößelmann 2017 | *P. aeruginosa* | 600 nm | Magnetic | A (external incubator) |
| Takahashi 2015, **Flexostat/Fluorostat** | *S. cerevisiae*, *E. coli* | 600 or 650 nm | Magnetic | A (external incubator) |
| Miller 2013, **Ministat array** | *S. cerevisiae* | - | Bubbling air | A (heating block) |
| Toprak 2012, **Morbidostat** (Toprak 2013) | *E. coli* | 950 nm | Magnetic | A (external incubator) |
| Marliere 2011 | *E. coli* | 880 nm | * | * |
| Tomson 2006, **Adaptastat** | *E. coli* | 600 nm | * | * |

**Table S1B - Complementary characteristics of already published continuous culture systems (continued).** [a] organisms are bacteria unless specified otherwise: *Bacillus subtilis*, *Subtilis Phage Pavia 1* (phage), *Escherichia coli*, *Euglena gracilis* (algae), *Lactococcus cremoris*, *Saccharomyces cerevisiae* (yeast), *Serratia marcescens*, *Pseudomonas aeruginosa*. [b] I (individually), A (all at once). * Information not available in publication, supplementary information or associated online data.

| Publication | Experiment control | Actuators and sensors management |
|---|---|---|
| This work, **ModuloStat** | Raspberry Pi; Perl, R | Espressif ESP32; C++ |
| Gopalakrishnan 2022, **EVE** | Raspberry; Python | |
| Pen 2021 | PC or Raspberry Pi; Python | Arduino Mega; C |
| Gervasi 2021 | Raspberry Pi; Node-RED | Espressif ESP32; C++ |
| Leyn 2021 | PC; MegunoLink | Arduino Mega; C |
| Steel 2020, **Chio.Bio** | BeagleBone Black; Python | |
| Ekkers 2020, **Omnistat** (Ekkers 2022) | PC; LabVIEW | Commercial complete devices |
| McGeachy 2019 | PC; Arduino IDE serial connection | Adafruit Feather M0; C++ |
| Guarino 2019 | Arduino Mega; * | |
| Wong 2018, **eVOLVER** (Heins 2019, García-Ruano 2023) | Raspberry Pi; Python | SAMD 21 Arduino; C |
| Hoffmann 2017 | PC; Python | Arduino Nano; C |
| Dößelmann 2017 | PC; Python | Arduino Mega; C |
| Takahashi 2015, **Flexostat/Fluorostat** | PC; Python | ATmega164; C |
| Miller 2013, **Ministat array** | None | Commercial complete devices |
| Toprak 2012, **Morbidostat** (Toprak 2013) | PC; Matlab | Commercial acquisition and control device |
| Marliere 2011 | *; * | *; * |
| Tomson 2006, **Adaptastat** | PC; BioXpert | *; * |

**Table S1C - Complementary characteristics of already published continuous culture systems (continued).** * Information not available in publication, supplementary information or associated online data.

| strain | genotype | reference |
|---|---|---|
| BSB1 | | Nicolas *et al.* 2012 |
| MS | *trpC2* Δ*ICEBs1*, Δ*PBSX*, Δ*SPB*, Δ*skin*, Δ(*gtaB yvzH ggaB ggaA yvzI yvzE*), Δ*upp*::[λpR-*neo*] | Dervyn *et al.* 2023 |
| $MS_{cat}$ | *trpC2* Δ*ICEBs1*, Δ*PBSX*, Δ*SPB*, Δ*skin*, Δ(*gtaB -yvzH ggaB ggaA yvzI yvzE*), Δ*upp*::[λpR-*cat*] | Dervyn *et al.* 2023 |
| del1 | *trpC2* Δ*ICEBs1*, Δ*PBSX*, Δ*SPB*, Δ*skin*, Δ(*gtaB -yvzH ggaB ggaA yvzI yvzE*), Δ*upp*::[λpR-*neo*], Δ*pro2* | lab collection |
| del2 | *trpC2* Δ*ICEBs1*, Δ*PBSX*, Δ*SPB*, Δ*skin*, Δ(*gtaB -yvzH ggaB ggaA yvzI yvzE*), Δ*upp*::[λpR-*neo*], Δ*pro2*, Δ*subti* | lab collection |
| del3 | *trpC2* Δ*ICEBs1*, Δ*PBSX*, Δ*SPB*, Δ*skin*, Δ(*gtaB -yvzH ggaB ggaA yvzI yvzE*), Δ*upp*::[λpR-*neo*], Δ*pro2*, Δ*subti*, Δ*pro1* | lab collection |
| del4 | *trpC2* Δ*ICEBs1*, Δ*PBSX*, Δ*SPB*, Δ*skin*, Δ(*gtaB -yvzH ggaB ggaA yvzI yvzE*), Δ*upp*::[λpR-*neo*], Δ*pro2*, Δ*subti*, Δ*pro1*, Δ*baci* | lab collection |
| del5 | *trpC2* Δ*ICEBs1*, Δ*PBSX*, Δ*SPB*, Δ*skin*, Δ(*gtaB -yvzH ggaB ggaA yvzI yvzE*), Δ*upp*::[λpR-*neo*], Δ*pro2*, Δ*subti*, Δ*pro1*, Δ*baci*, Δ*srf* | lab collection |
| del6 | *trpC2* Δ*ICEBs1*, Δ*PBSX*, Δ*SPB*, Δ*skin*, Δ(*gtaB -yvzH ggaB ggaA yvzI yvzE*), Δ*upp*::[λpR-*neo*], Δ*pro2*, Δ*subti*, Δ*pro1*, Δ*baci*, Δ*srf*, Δ*pro3* | lab collection |
| del7 | *trpC2* Δ*ICEBs1*, Δ*PBSX*, Δ*SPB*, Δ*skin*, Δ(*gtaB -yvzH ggaB ggaA yvzI yvzE*), Δ*upp*::[λpR-*neo*], Δ*pro2*, Δ*subti*, Δ*pro1*, Δ*baci*, Δ*srf*, Δ*pro3*, Δ*cory* | lab collection |
| del8 | *trpC2* Δ*ICEBs1*, Δ*PBSX*, Δ*SPB*, Δ*skin*, Δ(*gtaB -yvzH ggaB ggaA yvzI yvzE*), Δ*upp*::[λpR-*neo*], Δ*pro2*, Δ*subti*, Δ*pro1*, Δ*baci*, Δ*srf*, Δ*pro3*, Δ*cory*, Δ*basin* | lab collection |
| del9 | *trpC2* Δ*ICEBs1*, Δ*PBSX*, Δ*SPB*, Δ*skin*, Δ(*gtaB -yvzH ggaB ggaA yvzI yvzE*), Δ*upp*::[λpR-*neo*], Δ*pro2*, Δ*subti*, Δ*pro1*, Δ*baci*, Δ*srf*, Δ*pro3*, Δ*cory*, Δ*basin*, Δ*spo* | lab collection |
| del9trp+ | Δ*ICEBs1*, Δ*PBSX*, Δ*SPB*, Δ*skin*, Δ(*gtaB -yvzH ggaB ggaA yvzI yvzE*), Δ*upp*::[λpR-*neo*], Δ*pro2*, Δ*subti*, Δ*pro1*, Δ*baci*, Δ*srf*, Δ*pro3*, Δ*cory*, Δ*basin*, Δ*spo* | lab collection |
| IC (del10) | Δ*ICEBs1*, Δ*PBSX*, Δ*SPB*, Δ*skin*, Δ(*gtaB -yvzH ggaB ggaA yvzI yvzE*), Δ*upp*::[λpR-*cat*], Δ*pro2*, Δ*subti*, Δ*pro1*, Δ*baci*, Δ*srf*, Δ*pro3*, Δ*cory*, Δ*basin*, Δ*spo*, Δ*fla* | this work |
| del11 | Δ*ICEBs1*, Δ*PBSX*, Δ*SPB*, Δ*skin*, Δ(*gtaB -yvzH ggaB ggaA yvzI yvzE*), Δ*upp*::[λpR-*cat*], Δ*pro2*, Δ*subti*, Δ*pro1*, Δ*baci*, Δ*srf*, Δ*pro3*, Δ*cory*, Δ*basin*, Δ*spo*, Δfla, Δ*eps* | this work |
| del12 | Δ*ICEBs1*, Δ*PBSX*, Δ*SPB*, Δ*skin*, Δ(*gtaB -yvzH ggaB ggaA yvzI yvzE*), Δ*upp*::[λpR-*cat*], Δ*pro2*, Δ*subti*, Δ*pro1*, Δ*baci*, Δ*srf*, Δ*pro3*, Δ*cory*, Δ*basin*, Δ*spo*, Δfla, Δ*eps*, Δ*tasA* | this work |
| del13 | Δ*ICEBs1*, Δ*PBSX*, Δ*SPB*, Δ*skin*, Δ(*gtaB -yvzH ggaB ggaA yvzI yvzE*), Δ*upp*::[λpR-*cat*], Δ*pro2*, Δ*subti*, Δ*pro1*, Δ*baci*, Δ*srf*, Δ*pro3*, Δ*cory*, Δ*basin*, Δ*spo*, Δfla, Δ*eps*, Δ*tasA*, Δ*bslA* | this work |
| del14 | Δ*ICEBs1*, Δ*PBSX*, Δ*SPB*, Δ*skin*, Δ(*gtaB -yvzH ggaB ggaA yvzI yvzE*), Δ*upp*::[λpR-*cat*], Δ*pro2*, Δ*subti*, Δ*pro1*, Δ*baci*, Δ*srf*, Δ*pro3*, Δ*cory*, Δ*basin*, Δ*spo*, Δfla, Δ*eps*, Δ*tasA*, Δ*bslA*, Δ*pgs* | this work |
| ZB (del15) | Δ*ICEBs1*, Δ*PBSX*, Δ*SPB*, Δ*skin*, Δ(*gtaB -yvzH ggaB ggaA yvzI yvzE*), Δ*upp*::[λpR-*cat*], Δ*pro2*, Δ*subti*, Δ*pro1*, Δ*baci*, Δ*srf*, Δ*pro3*, Δ*cory*, Δ*basin*, Δ*spo*, Δfla, Δ*eps*, Δ*tasA*, Δ*bslA*, Δ*pgs*, Δ*yda* | this work |

**Table S2**: ***B. subtilis*** **zero-biofilm (ZB) genetic construction** with complete genotype of all ancestral strains from the BSB1 strain reference genome (Genbank: AL009126).

| Strain name | Last deletion | Gene (from:to) | Position (from:to) | Rationale |
|---|---|---|---|---|
| del1 | Δ*pro2* | *ydcL:yddM* | 529406:549932 | prophages: no lysis of stressed cells |
| del2 | Δ*subti* | *sboX:albG* | 3836106:3842995 | antibacterial activity: no negative effect in the event of population mixing |
| del3 | Δ*pro1* | *alkA:skfH* | 202551:220032 | prophages: no lysis of stressed cells |
| del4 | Δ*baci* | *ytpB:ytoA* | 3121595:3124007 | antibacterial activity: no negative effect in the event of population mixing |
| del5 | Δ*srf* | *sfrAB:sfp* | 391044:408135 | antibacterial activity: no negative effect in the event of population mixing |
| del6 | Δ*pro3* | *ydzT:ydjC* | 652043:664671 | prophages: no lysis of stressed cells |
| del7 | Δ*cory* | *dhbF:dhbA* | 3280719:3292296 | antibacterial activity: no negative effect in the event of population mixing |
| del8 | Δ*basin* | *ytpB:ytoA* | 3868287:3874180 | antibacterial activity: no negative effect in the event of population mixing |
| del9 | Δ*spo* | *spoIIAA:sigF* | 2443421:2445002 | sporulation: no spore able to resist stress |
| IC (del10) | Δ*fla* | *flgB:ylxH* | 1691278:1711685 | motility: to limit potential swimming back in silicone tubing |
| del11 | Δ*eps* | *epsO:epsA* | 354185:3529940 | exopolysaccharide biosynthesis * |
| del12 | Δ*tasA* | *tasA* | 2553082:2553866 | major component of biofilm matrix, forms bundles of fibres * |
| del13 | Δ*bslA* | *bslA* | 3187526:31888043 | bacterial hydrophobin, forms water-repellent surface layer of the biofilm |
| del14 | Δ*pgs* | *pgsB:pgsC* | 3698999:3700606 | poly-gamma-glutamate biosynthesis |
| ZB (del15) | Δ*yda* | *ydaJ:ydajN* | 479009:485950 | extracellular polysaccharide synthesis and transport * |

**Table S3**: ***B. subtilis* zero-biofilm (ZB) strain deleted regions** relative to BSB1 reference genome (Genbank: AL009126). * mutated regions sequenced in formed biofilm during experiments with intermediate lineages.

### SUPPLEMENTARY FIGURES S1-S14

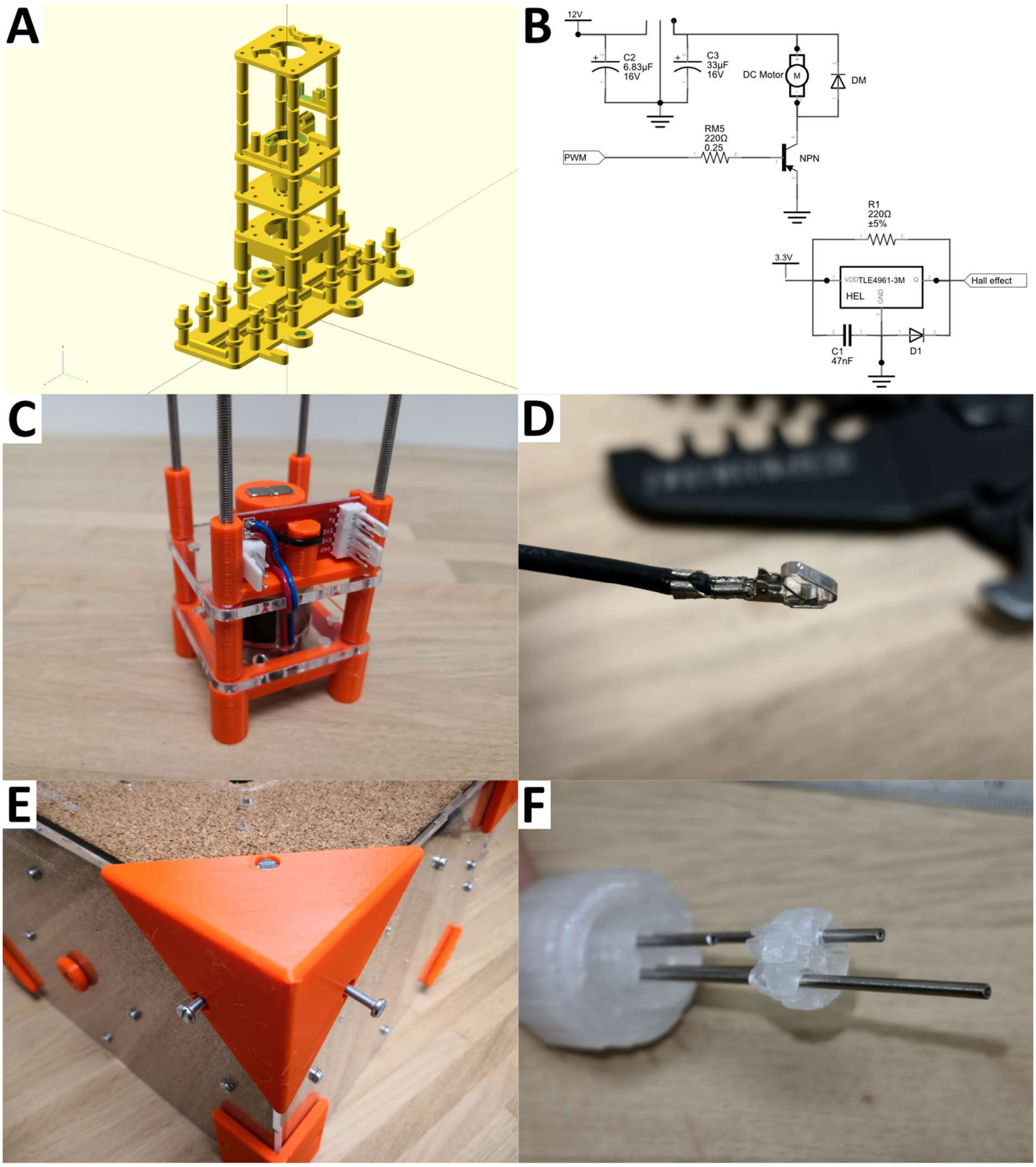

**Figure S1: Illustrative pictures of the ModuloStat documentation and assembly guide.** Various pictures were sampled from the online documentation illustrating the system and its assembly steps. **A** Openscad 3D rendering of the mini-bioreactor holder. **B** Electronic circuits schema of the hall effect sensor retro-controlled magnetic stirring. **C** Intermediate step of the mini-bioreactor assembly for magnetic stirring electronics. **D** Focus on the connectors clamping method. **E** Intermediate step of the box assembly (3D-printed corner installation on PMMA and cork sides with screws). **F** Intermediate step of the mini-bioreactor screw-cap assembly for friction loving device installation on stainless-steel tubing.

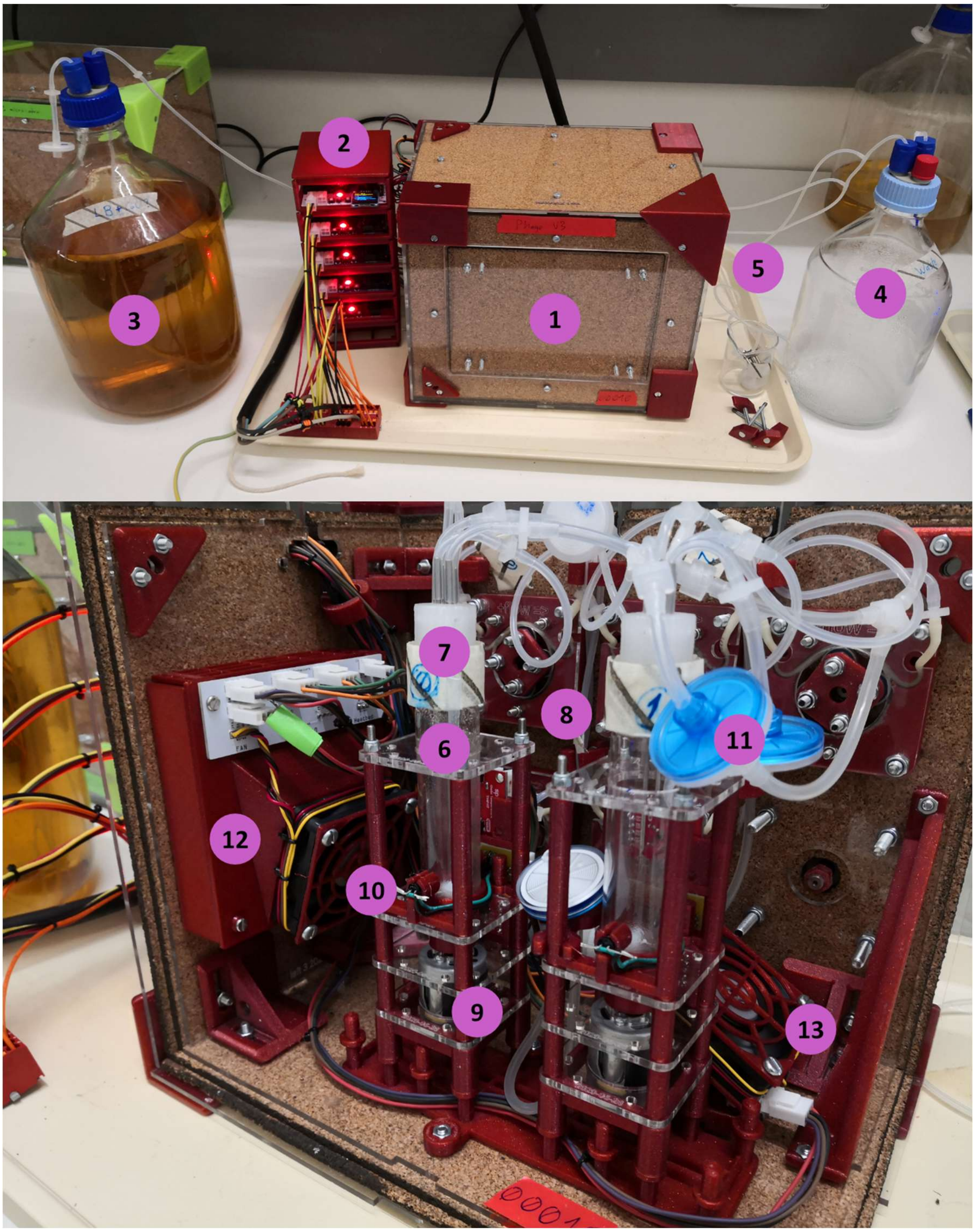

**Figure S2: Composition of the ModuloStat system** with (**1**) thermoregulated box made of laser-cut PMMA and cork (closed view), (**2**) stack of electronic circuits with one ESP32 per layer, (**3**) feeding bottle (5 L here), (**4**) waste bottle (overflow), (**5**) sampling port derivation system preserving bioreactor's sterility, (**6**) bioreactor from a commercial borosilicate tube (Sheaton 24 mL type), (**7**) 3D-printed polypropylene bioreactor screwcap sealed with laser-cut silicone sheet, (**8**) 3D-printed peristaltic pump head, (**9**) magnetic stirring with 3D-printed head, (**10**) coupled light source (LED with emission peak at 940 nm) and sensor (phototransistor) for optical density measure, (**11**) bioreactors aeration through a silicone tube circuit (3 mm internal diameter) and air filters (0.2 µm), (**12**) heating system made from a heat plate and a fan, and (**13**) secondary fan to homogenise temperature inside the box.

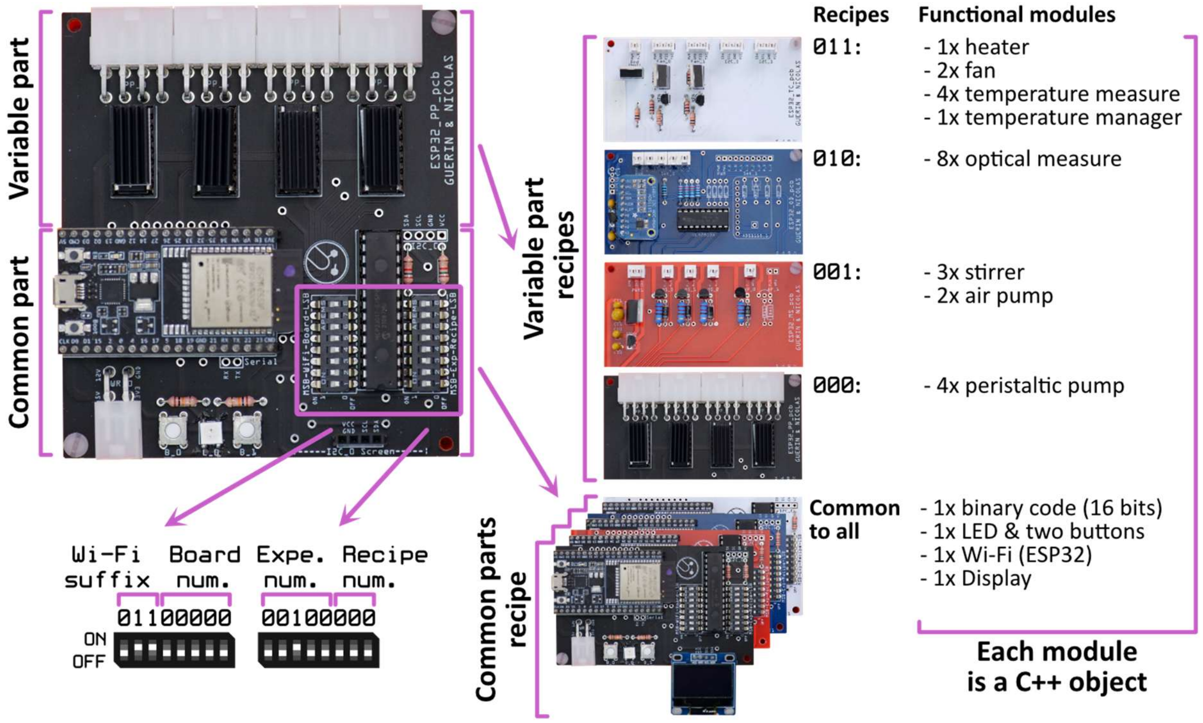

**Figure S3: Modular design of ModuloStat board electronic circuits.** The board contains several electronic functional modules, some are common to all boards (common part, 89 x 47 mm) and some are specific to each type of board (variable part, 89 x 53 mm). Each functional module is programmatically managed on the ESP32 micro-controller as a C++ object dynamically instantiated at boot according to recipe number. The recipe number is determined using 3 digits of the 2x8 bits binary encoders present on the common part. The upper left of the figure shows a single PCB board dedicated to peristaltic pump control. The lower left of the figure explains the use of the 2x8 binary encoders. The center of the figure represents the different common and variable parts of the PCBs for the four types of ModuloStat boards used in this work. The right of the figure lists the modules included in each type of board recipe.

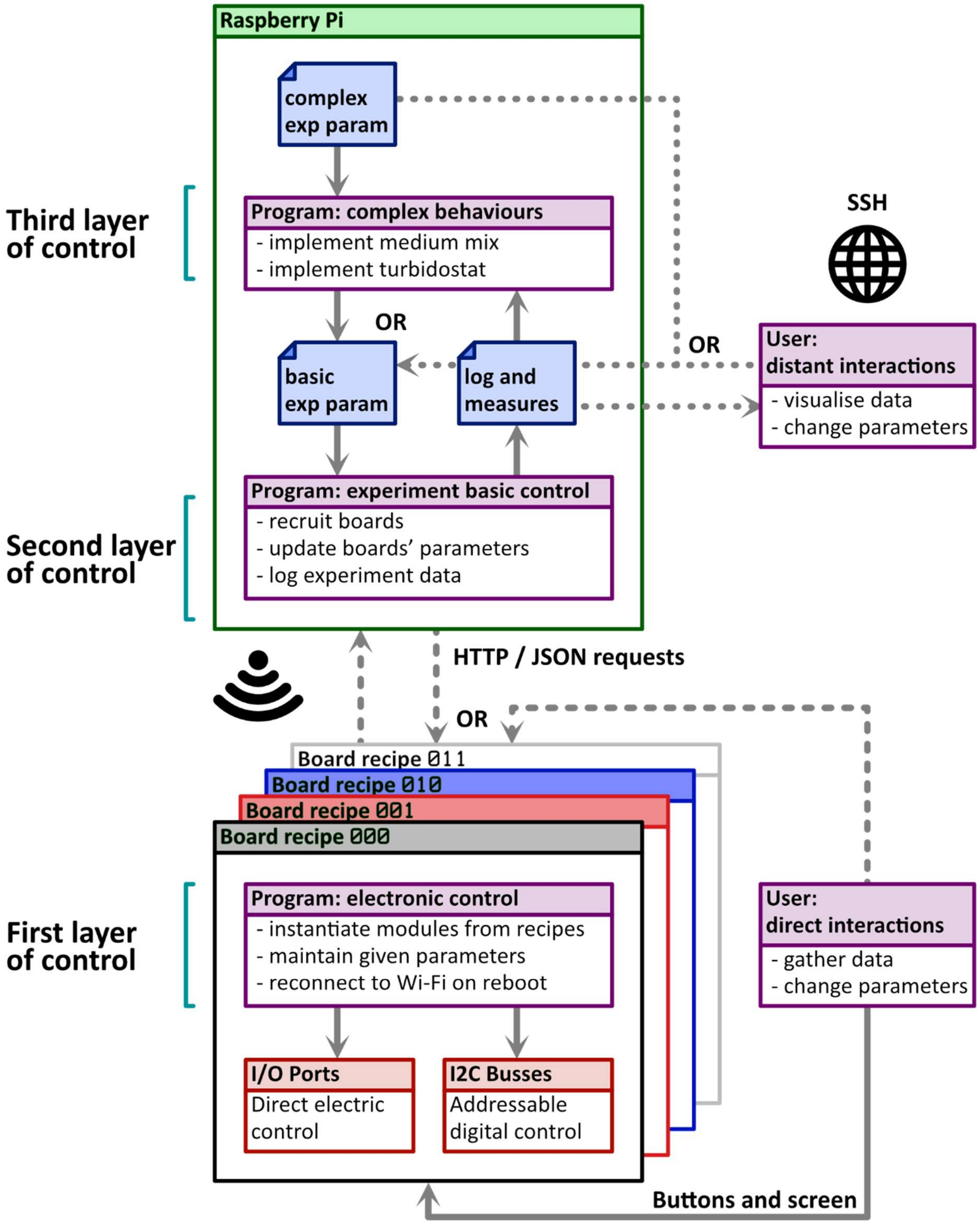

**Figure S4**: **Schematic representation of the different layers of control of an experiment.** Solid arrows: Read/write files or electronics, read board screen. Dashed arrows: HTTP/JSON interactions (Wi-Fi) with arrows indicating the initial requests direction. Dotted arrows: SSH interactions for parameter files editions or data gathering.

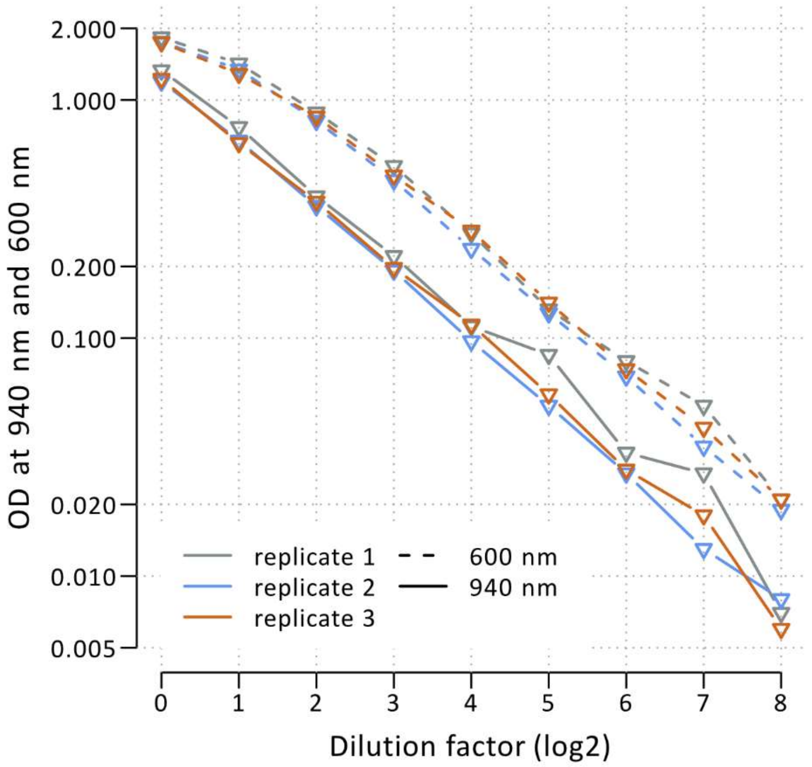

**Figure S5: Comparison of OD values measured at 600 nm and 940 nm.** Saturated cultures of *B. subtilis* grown in LB were mixed with 1mg/mL chloramphenicol to stop cell division. LB medium supplemented with 1mg/mL chloramphenicol were used for dilution. Successively, OD at 600 nm and 940 nm were measured with a UV-1900 Shimadzu spectrophotometer, and two fold dilution was then performed by adding equal amounts of the LB medium supplemented with chloramphenicol. OD blanks references were performed using LB medium supplemented with chloramphenicol. The near infrared OD (940 nm) performs almost linearly, even at the highest concentrations of bacteria.

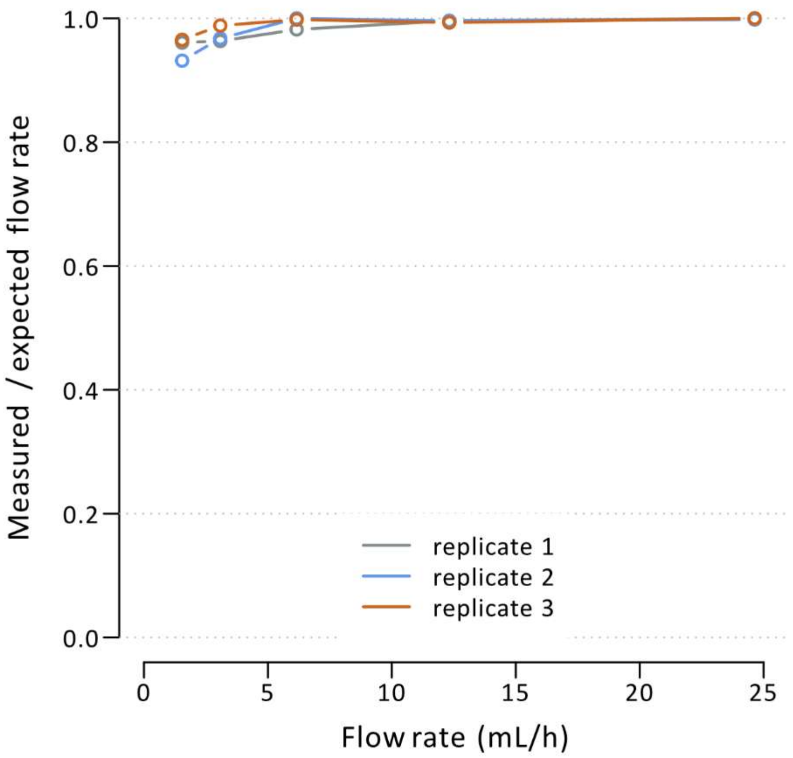

**Figure S6: Deviation between measured and expected flow rates at very low pump speed.** The expected flow rate is calculated from the speed imposed to the peristaltic pump using the conversion factor of 4.93 ml/h for one rpm. This is the same data presented in Figure 3B, but in a different form.

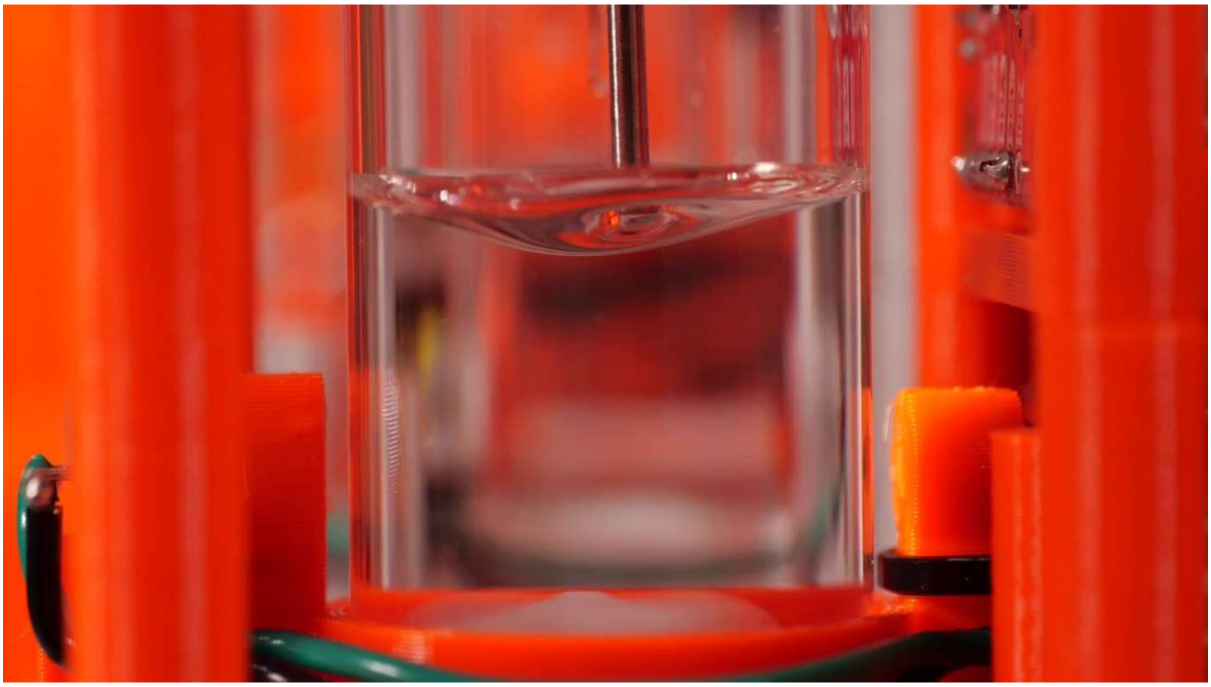

**Figure S7: Setting the volume of the culture by adjusting the height of the outflow tubing mouth.** The liquid surface tension between the vortex and the stainless-steel tubing introduces a small uncertainty in the volume of the culture.

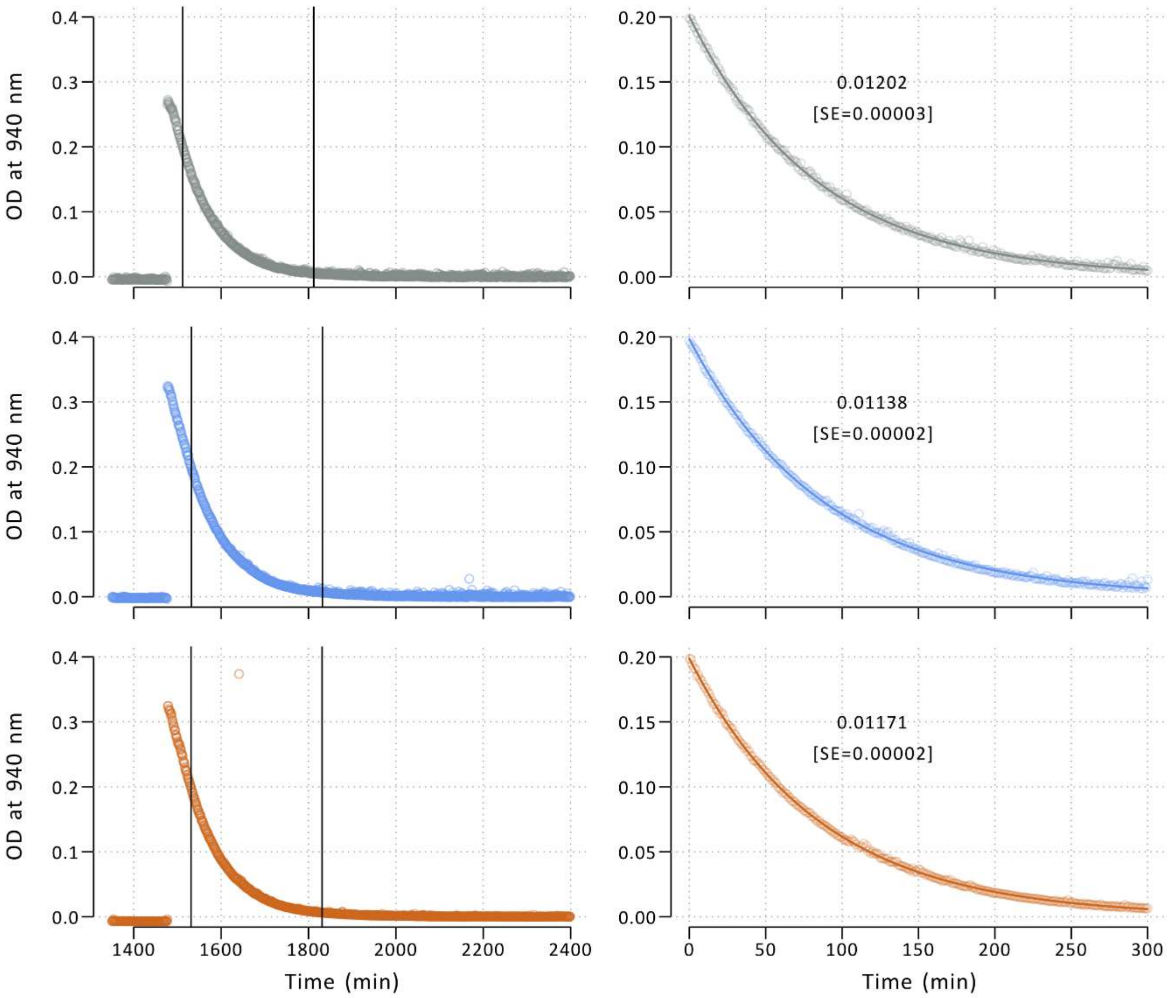

**Figure S8: Dilution rate characterisation from non-growing *B. subtilis* cells continuous dilution.** Saturated cultures of *B. subtilis* grown in LB medium were mixed with 1mg/mL chloramphenicol to stop cell division. Non-growing saturated *B. subtilis* cultures were pipetted in the emptied mini-bioreactors of the assembled ModuloStat with feeding and outflow pumps turned off. In the closed box when target temperature was reached, the flow rate of the continuous dilution with LB supplemented with chloramphenicol were set to 0.0855 $mL.min^{-1}$ (1.04 rpm according to the correspondence coefficient of 4.93 $mL.h^{-1}$ for 1 rpm established using the data of Figure 3B) and *in situ* OD were measured every minute at 940 nm. OD blank references were performed using LB medium supplemented with chloramphenicol at the end of the dilution process. Given the culture volume of 7 mL, this flow rate corresponded to a target dilution rate of 0.0122 $min^{-1}$. The dilution rates were estimated with the "nls" function of the R package "stats" in the three bioreactors on the data points collected during the 5 hours after crossing down the OD threshold of 0.2 (subset of data between the vertical bars in the left panel and represented in the right panel with the fitted dilution model shown in green). The three estimated values and standard errors are reported in the right-panel plots (mean 0.117 $min^{-1}$, coefficient of variation 2.7%).

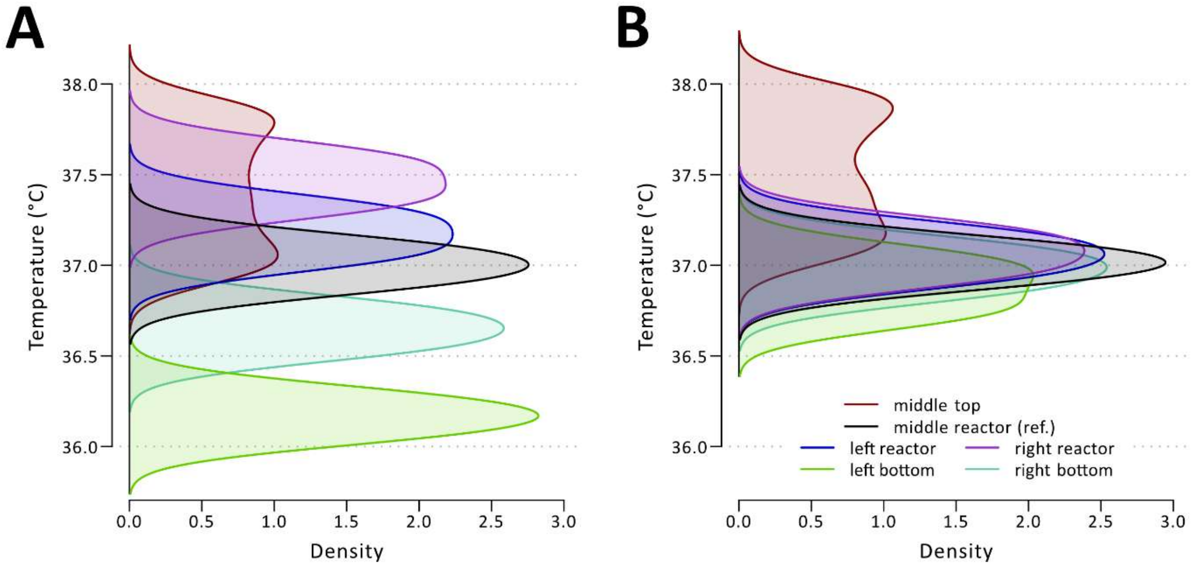

**Figure S9: Temperature distribution in the thermoregulated box with one or two fans.** Density distribution of temperature measured simultaneously at six different points inside the box over a period of three hours. The target temperature was set to 37°C. Densities were calculated using the R function "density" with a bandwidth of 0.1. **A** Distribution of temperatures when a single fan (main fan) is installed in the box directly next to the heater (left of the box, see Figure S2). **B** Distribution of temperatures when a secondary fan is installed at the bottom right of the box (see Figure S2). Same data as in Figure 3C.

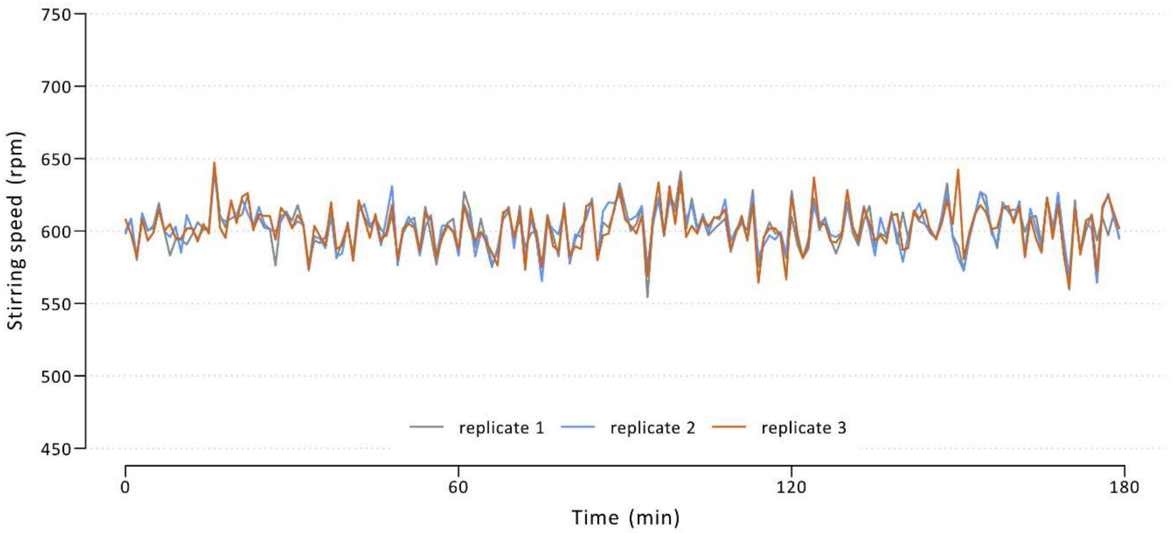

**Figure S10: Control of the magnetic stirring speed.** Speed is measured using a Hall effect sensor over a period of three hours. The input current of each stirring motor is adjusted (retro-controlled) based on this measured value through pulse width modulation (PWM). For a target steering speed of 600 rpm, the mean of the stirring speed measured was 602.5 rpm with a standard deviation of 14.3. Represented speed measures are sampled every minute.

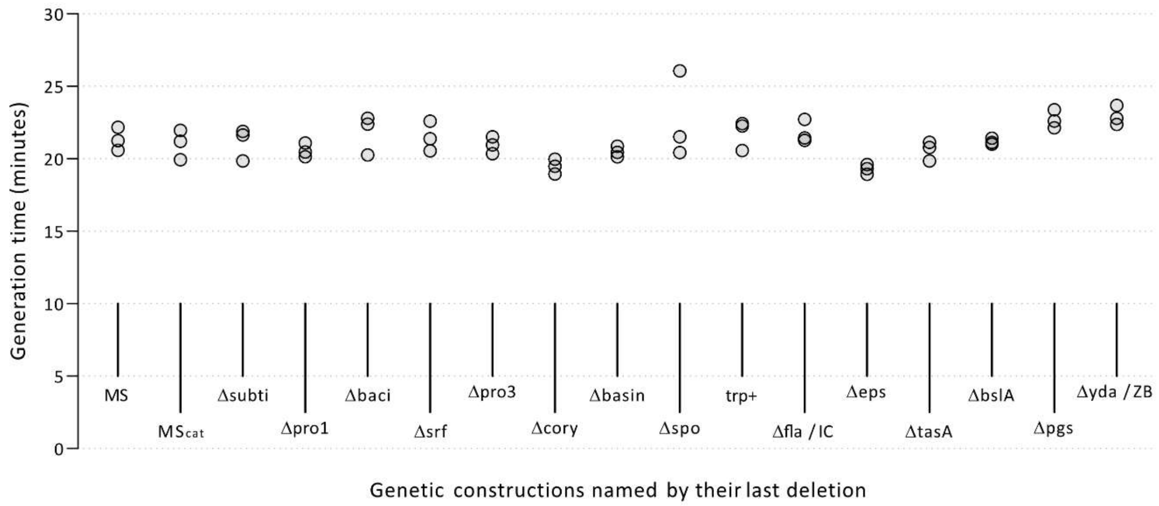

**Figure S11**: **Generation times in the construction lineage of the *B. subtilis* ZB strain.** Strains are represented from left to right in the order of the construction lineage of the ZB strain (Figure 4C and Table S2). Strains are named by their last genetic modification (Table S3). Exponential-phase cultures of each strain were diluted 400 fold in 96-well plates (Greiner) in a final volume of 100 µL of LB medium. The plate was grown at 37°C with agitation, and OD at 600 nm measured every 10 minutes (Synergy HTX). The generation time was estimated using a linear regression ($R^2 \geq 0.98$) over the logarithm of 6 OD measurements in the exponential growth phase.

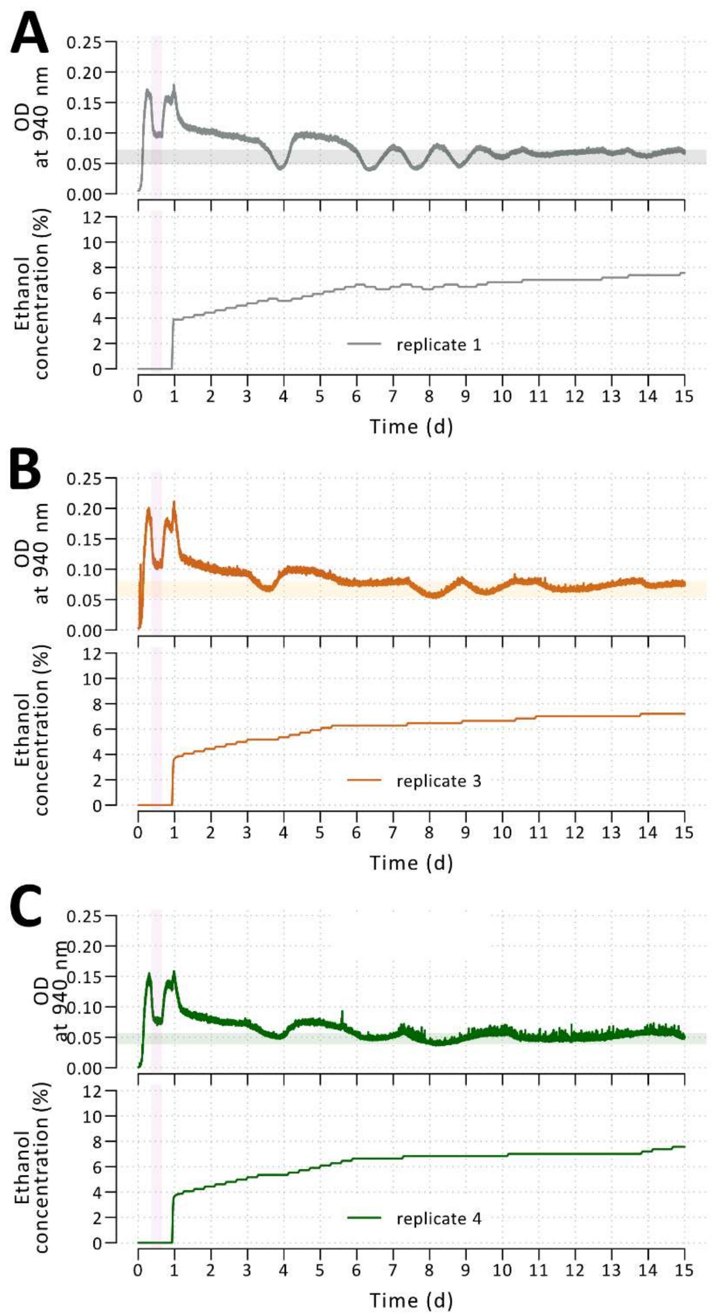

**Figure S12: Trajectories of cultures during the evolution of resistance to ethanol stress.** The same representation as in Figure 6B, but for the other three replicates.

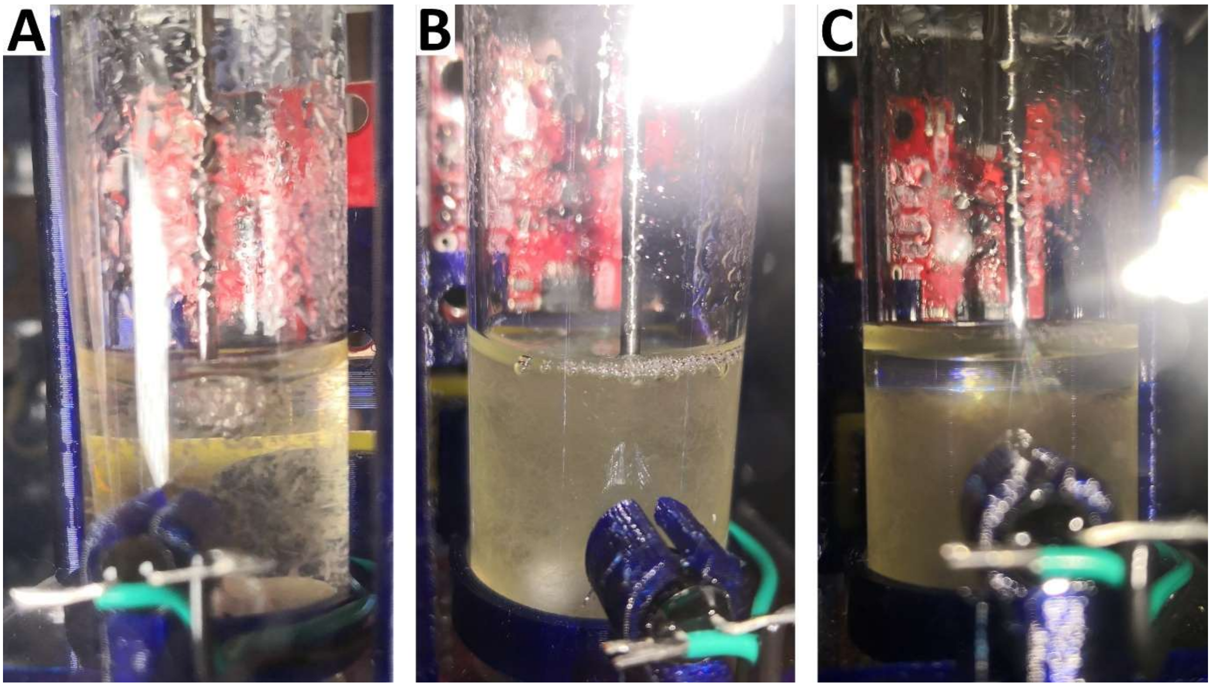

**Figure S13: Images of the "flake" phenotype that appeared during the evolution of resistance to ethanol stress. A** The culture of replicate 1 at 23 days of culture when aggregates ("flakes") started to accumulate. The image was taken under standard stirring at 600 rpm. **B** The same culture at 41 days of culture, i.e. after 7 days in the regime with ethanol concentration fixed at 7.5%. **C** The same as **B** but after 10 minutes without stirring; the "flakes" have sedimented.

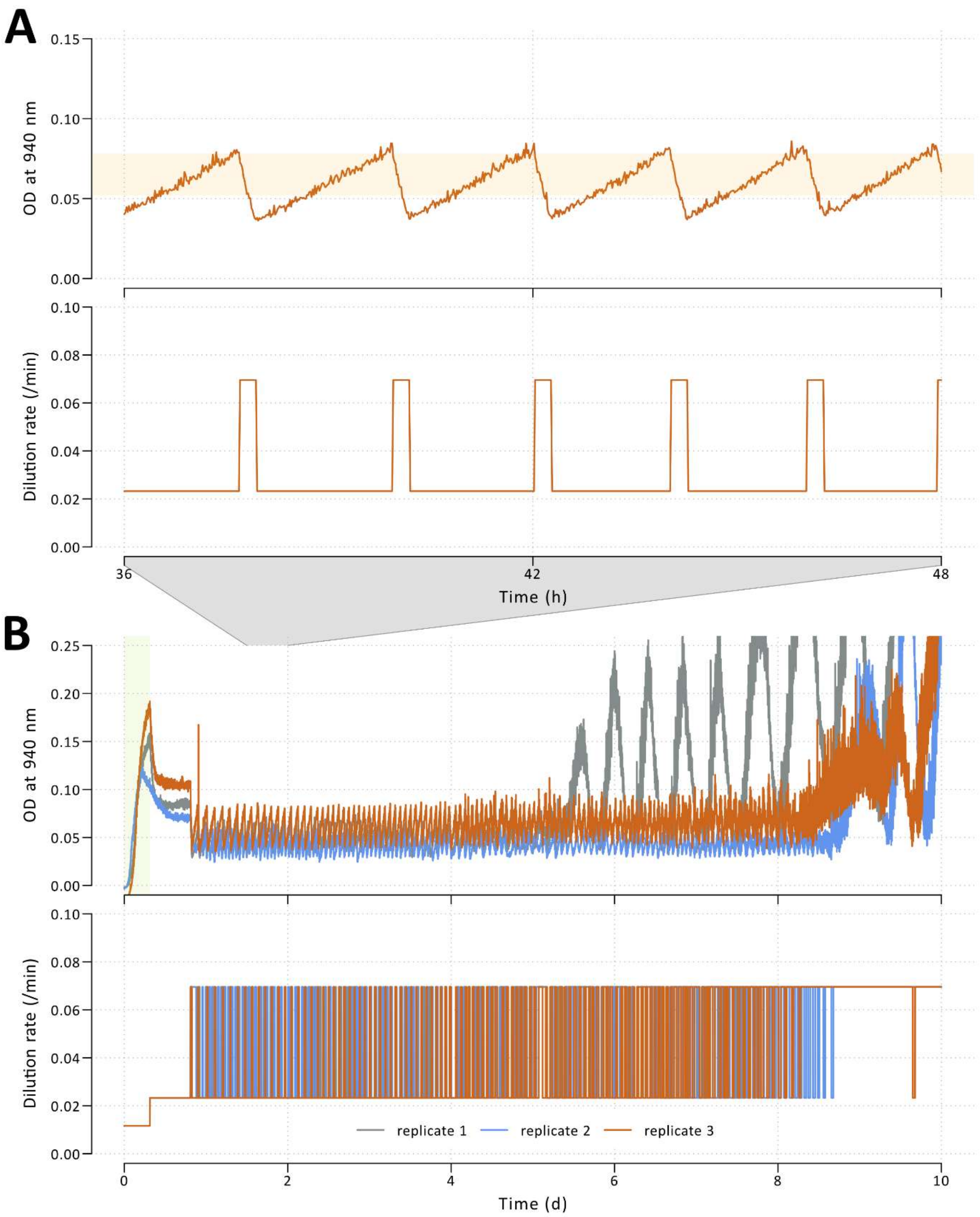

**Figure S14: Trajectories of turbidostat cultures of the ZB strain.** Two dilution rates are alternatively applied after the onset of the culture: the low dilution rate corresponds to a steady-state generation time of 30 minutes; the high dilution rate is three times higher and would thus correspond to a steady-state generation time of 10 minutes (which cannot be sustained by *B. subtilis*). The switch between the two dilution rates is applied when OD is above or below two OD target thresholds which are set to 50% and 75% of the OD at steady state of the low dilution rate. **A** OD trajectory and dilution rate over one day of culture. The OD target thresholds delimit the light-orange horizontal in the OD plot. **B** Trajectories for the three replicates over the entire experiment. Of note, we noticed that the outflow silicone tubing of replicate 2 was pinched during the beginning of the experiment, resulting in an inaccurate estimation of the optical density at 30 minutes of generation time and probable more stringent OD thresholds. After five to nine days, aggregates appeared in each culture, resulting in a noisier optical density (OD) that did not respond well to changes in dilution rates probably due to a trend toward sedimentation.
